# Dynamics of adaptation in an anisotropic phenotype-fitness landscape

**DOI:** 10.1101/623330

**Authors:** F. Hamel, F. Lavigne, G. Martin, L. Roques

## Abstract

We study the dynamics of adaptation of a large asexual population in a *n*-dimensional phenotypic space, under anisotropic mutation and selection effects. When *n* = 1 or under isotropy assumptions, the ‘replicator-mutator’ equation is a standard model to describe these dynamics. However, the *n*-dimensional anisotropic case remained largely unexplored.

We prove here that the equation admits a unique solution, which is interpreted as the phenotype distribution, and we propose a new and general framework to the study of the quantitative behavior of this solution. Our method builds upon a degenerate nonlocal parabolic equation satisfied by the distribution of the ‘fitness components’, and a nonlocal transport equation satisfied by the cumulant generating function of the joint distribution of these components. This last equation can be solved analytically and we then get a general formula for the trajectory of the mean fitness and all higher cumulants of the fitness distribution, over time. Such mean fitness trajectory is the typical outcome of empirical studies of adaptation by experimental evolution, and can thus be compared to empirical data.

In sharp contrast with the known results based on isotropic models, our results show that the trajectory of mean fitness may exhibit (*n* − 1) plateaus before it converges. It may thus appear ‘non-saturating’ for a transient but possibly long time, even though a phenotypic optimum exists. To illustrate the empirical relevance of these results, we show that the anisotropic model leads to a very good fit of *Escherichia coli* long-term evolution experiment, one of the most famous experimental dataset in experimental evolution. The two ‘evolutionary epochs’ that have been observed in this experiment have long puzzled the community: we propose that the pattern may simply stem form a climbing hill process, but in an anisotropic fitness landscape.

## 1 Introduction

### Biological motivation

Understanding the adaptation of asexual organisms (such as viruses, bacteria or cancer cells) under the combined effects of selection and mutation is a fundamental issue in population genetics.

In parallel, the development of experimental evolution (especially in microbes) has made it possible to compare observed dynamics, in the lab, with alternative models or infer various model parameters from the data (for a recent special issue on this subject see [38]). Still, the problem of modelling asexual evolutionary dynamics is inherently complex (discussed in *e.g*. [16]): recurrent mutation changes the key parameters of the dynamical system constantly, competition between numerous, ever changing types must be described, and both mutational and demographic (birth/death) events are stochastic in nature.

Recent models of asexual adaptation seek to follow the dynamics of the full distribution of fitness - the expected reproductive output of a lineage - within populations. Contrarily to other approaches, such as ‘origin fixation models’ which only follow the expected mean fitness 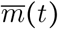 [40], these models do not make a low polymorphism assumption, but in exchange for ignoring or simplifying the stochastic components of the dynamics. The resulting outcome is a partial differential equation (PDE) or an integro-differential equation (IDE), that typically describes the dynamics of the distribution of a single ‘trait’. In some cases, this trait may be fitness itself as in [1, 17, 42], leading to equations of the form:

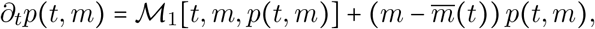

where the variable *m* ∈ ℝ is the fitness and 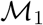 is a differential or an integral operator describing the effect of mutations on the distribution *p*(*t*, ·) of fitness. Other models describe the distribution of a given trait *x* ∈ ℝ determining fitness (birth rate, phenotype), as in [2, 8]:

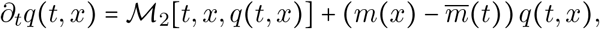

where *m*(*x*) is a function which describes the relationship between the trait *x* and the fitness. Here, 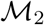 is a differential or an integral operator describing the effect of mutations on the distribution *q*(*t*, ·) of the trait *x*. In these two equations, the last term 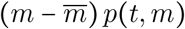 (resp. 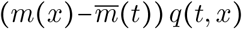) corresponds to the effects of selection and will be explained later. Broadly, it emerges when the growth rate *m*(*x*) (in the absence of competition) of a genotype with phenotype *x* depends only on its own phenotype. This corresponds to the classic frequency-independent model of selection.

One established finding from empirical fitness trajectories is that epistasis, namely the fact that the distribution of a mutation’s effect on fitness depends on its ancestor’s genetic background, must be accounted for to explain the data (*e.g*. [21]). More precisely, fitness trajectories tend to decelerate over time, implying *a priori* that beneficial mutations become weaker and/or rarer as the population adapts (ignoring deleterious ones, which is of course debatable). The question is then: which particular form of epistasis does explain/predict the data best? A common metaphor to explain this decelerating pattern in fitness trajectories is to invoke some form of ‘fitness landscape’ connecting genotypes or phenotypes with fitness, as in the function *m*(*x*) above, with one or several adaptive peak(s) where fitness is maximal (discussed in [11]).

In this view, deceleration in fitness trajectories stems from the hill climbing process of adaptation up a fitness peak. This view is appealing because of its intuitive/visual illustration, but also because of a particular form of landscape, Fisher’s geometrical model (FGM; a single peak phenotype-fitness landscape), has shown promising potential when compared to empirical measures of fitness epistasis. In the FGM, a multivariate phenotype at a set of *n* traits (a vector **x** ∈ ℝ^*n*^) determines fitness. The most widely used version assumes a quadratic form of the Malthusian fitness function *m*(**x**), which decreases away from a single optimum **O** ∈ ℝ^*n*^[31, 41]. Without loss of generality, the traits can be defined such that the optimum sets the origin of phenotype space:

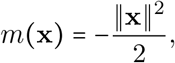

with ‖·‖ the Euclidian norm in ℝ^*n*^. To describe the mutation effects on phenotypes, the standard ‘isotropic Gaussian FGM’ uses a normal distribution 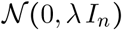 with λ > 0 the phenotypic mutational variance at each trait and *I_n_* the identity matrix.

Given the potential complexity of the relationship between genotype, phenotype and fitness, the FGM may appear highly oversimplified. However, several arguments suggest that it can be more than a mere heuristic and may be a relevant, empirically sound null model of adaptation (also discussed in a recent review of the FGM [41]). First [27], the Gaussian FGM can be retrieved from a much less constrained model of phenotype-to-fitness landscape: Random Matrix theory arguments show that it emerges as a limit of highly integrative phenotypic networks, near an optimum (with an arbitrary fitness function and distribution of the original phenotypes). Second, several assumptions of the classic FGM have been tested quite extensively on empirical distributions of mutation effects on fitness, showing its power to fit or predict the observed patterns. For example, the FGM has shown to accurately predict observed distributions of epistasis among random and beneficial mutations in a virus and a bacterium [28], to accurately fit patterns of epistasis among beneficial mutations in a fungus [39] or the pattern of re-adaptation (compensation) from different deleterious mutant backgrounds in a ‘recovery’ experiment with a bacterium [34]. It also accurately predicted how stressful conditions (lowering the parental fitness) affect the mean and variance of mutation fitness effects [30]. The assumption of a quadratic fitness function is perhaps the most critical to the patterns tested so far: various empirical tests showed that a deviation from the quadratic would actually lead to a lower fit of the data (see [13] and fig. 5 in [30]). Overall, this suggests that, in spite of its apparent simplicity, the FGM with a quadratic fitness function is not so oversimplified: it is a natural limit of a diversity of more complex models and is consistent with various empirical patterns of mutation fitness effects.

On the other hand, this ‘fitness peak’ view is challenged by the observation [45] that, in the longest evolution experiment ever undertaken (Long Term Evolution Experiment LTEE, in the bacterium *Escherishia coli*), fitness has not reached any maximum after more than 70,000 generations. It has been suggested [19] that this experiment actually shows a ‘two epochs’ dynamics with an initial (potentially saturating) fitness trajectory and a later and distinct nonsaturating dynamics. A similar pattern could be invoked in another mid-term experiment with an RNA virus [33]. Several other experiments did seem to yield a ‘true’ saturation (plateau) in fitness trajectories over several thousands of generations in *E. coli* (*e.g*. [9, 12, 22]), but they may simply be on a too short timescale to identify subsequent fitness increases. In fact, the LTEE itself did seem to show a strongly saturating pattern after 10,000 generations [24].

Overall, these different insights on epistasis and adaptation trajectories are difficult to reconcile under a single theory: why would a single peak model show a good fit to mutation epistasis over single mutations, or short term fitness trajectories, yet be invalid over the longer term? Is there really a two ‘epochs’ dynamics in some long-term trajectories, when and why? A desirable model should reconcile both timescales, describe both epistasis among random single mutations and in long term fitness trajectories. Proposed models that do accurately fit non-saturating trajectories [45] have the drawback that they only focus on beneficial mutations and do not yield any prediction on the distribution of epistasis among random mutations. On the other hand, simple fitness landscapes like the FGM do predict some short term patterns and the full distribution of epistasis among random mutants, but cannot yield a never-ending fitness trajectory (a fitness plateau must be reached at the optimum).

### Aim of the paper

In this article, we explore the possibility that an extended version of the FGM be able to fit multiple epochs fitness trajectories. The central extension proposed here is to consider anisotropy in the landscape, in that different underlying traits affect fitness differently and mutate differently. Namely, the fitness function associated with a trait **x** = (*x*_1_,…,*x_n_*) is:

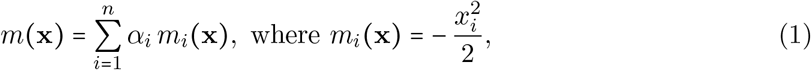

where the coefficients *α_i_* are positive. The mutation effects on phenotypes are described with an anisotropic Gaussian distribution 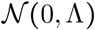, where Λ = diag (λ_1_,…,λ_*n*_) is any given positive diagonal matrix.

Using central limit and random matrix theory arguments [27], the FGM can be obtained as a limit of a much less constrained model where high-dimensional phenotypes integrate into a phenotypic network to a smaller set of traits that directly determine fitness, with an optimum. The resulting FGM, however, is not necessarily isotropic, it can also show a single dominant direction in phenotype space (that affects fitness much more) with all other directions remaining approximately equivalent. Our initial intuition is that, in these conditions, adaptation along the dominant direction will drive the early dynamics while a second ‘epoch’ will be visible when adaptation only proceeds along the remaining dimensions. Therefore, it seems natural to explore how such a model would fare when compared to the fitness trajectories of the LTEE. Yet, this requires deriving the fitness dynamics resulting from such a complex mutational model, in the presence of mutation and competition between asexual lineages. This is the aim of the present paper.

The existence of a phenotype optimum has been taken into account in recent PDE and IDE approaches (second class of models alluded to above). For instance, in [2, 3], the fitness depends on a single phenotypic trait *x* ∈ ℝ, through a function *m*(*x*) which admits an optimum. Extending these works to take into account the dependence of the fitness on several traits appears as a natural question; especially since we know that the number of traits affected by mutation and subject to selection critically affects the evolutionary dynamics [32, 44]. So far, and to the best of our knowledge, such mathematical models that take into account *n*-dimensional phenotypes together with the existence of a phenotype optimum always assume an isotropic dependence between the traits **x** ∈ ℝ^*n*^ and the fitness, and an isotropic effect of mutations on phenotypes (see [18] for an IDE approach and [32] for an approach based on the analysis of a PDE satisfied by a moment generating function of the fitness distribution). Our goal here is to extend these approaches in order to characterize the dynamics of adaptation in a *n*-dimensional phenotypic space, without making an isotropy assumption. For the sake of simplicity here, we ignore any stochastic component of the dynamics, but note that the same equations can be obtained explicitly as a deterministic limit of a stochastic model of mutation and growth [32].

### Model assumptions and definitions

We assume that a phenotype is a set of *n* biological traits, which is represented by a vector **x** ∈ ℝ^*n*^. To describe the evolution of the phenotypic distribution we begin with an intuitive description of the concept of fitness. In an asexual population of *K* types of phenotypes **x**_1_,…,**x**_*K*_, with *r*_max_ > 0 the growth rate of the optimal phenotype, the *Malthusian fitness, relative to the optimal phenotype* is denoted *m_j_* = *m*(**x**_*j*_) ≤ 0 for a phenotype **x**_*j*_ ∈ ℝ^*n*^. It is defined by the following relation: 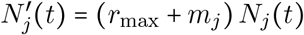, with *N_j_*(*t*) the abundance of the phenotype **x**_*j*_ at time *t*. When we sum these equations over all indexes *j* = 1,…,*K*, we obtain an ordinary differential equation for the total population size 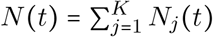 at time 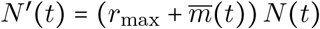, where the quantity 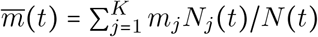 is the mean relative fitness in the population at time *t*. Now if we turn to the distribution of the phenotype frequencies *q*(*t*, **x**_*j*_) = *N_j_*(*t*)/*N*(*t*), we get the partial differential equation:

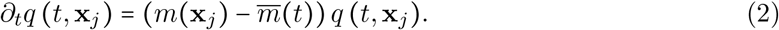

This equation can be generalized to a continuous distribution of phenotype frequencies (see *e.g*. [42]), as we assume in the sequel, with:

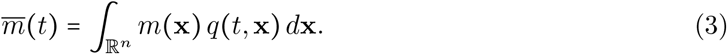

If 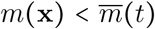, then (2) implies that the frequency of the phenotype **x** decreases, whereas if 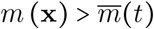, then the frequency increases.

We recall that mutation’s effects on phenotypes are assumed to follow an anisotropic Gaussian distribution 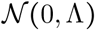, with Λ = diag(λ_1_,…,λ_*n*_) a positive diagonal matrix. Assuming a small mutational variance max λ_*i*_ << 1, and a mutation rate *U*, these mutational effects can be approximated by an elliptic operator 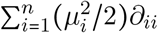, where 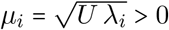 and *∂_ii_* denotes the second order partial derivative with respect to the *i*-th coordinate of **x** (or later **m** as in (17) below). We refer to Appendix A for further details on the derivation of this diffusion approximation. Intuitively, the regime where it applies is the ‘Weak Selection Strong Mutation’ (WSSM) regime (high rate of small effect mutations) where a wide diversity of lineages accumulate mutations and co-segregate at all times. The extreme alternative is the Strong Selection Weak Mutation (SSWM) regime where few mutations of large effect invade successively and the population is effectively close to monomorphic at all times. This latter regime will not be treated here.

Overall, the corresponding PDE describing the dynamics of the phenotype distribution *q*(*t*, **x**), under the combined effects of selection and mutation, is:

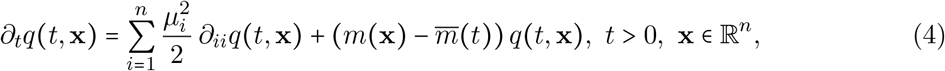

with 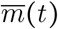 defined by (3). Recent studies [2] have already treated the case *n* = 1 (see also [3] for more general fitness functions). In this paper, we consider the general case *n* ≥ 1. Without loss of generality, we may assume in the sequel that *α*_1_ = … = *α*_*n*_ = 1 in (5), up to a scaling of the phenotype space 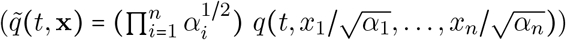. This leads to:

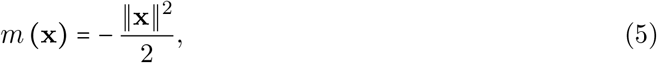

with ‖·‖ the standard Euclidian norm in ℝ^*n*^. Similarly, we could remove the anisotropy in the mutation effects, up to an other scaling of the phenotype space (defined this time by 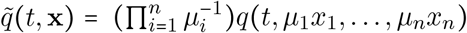), but in this case the coefficients *α_i_* would be replaced by 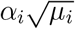 and hence cannot be taken all equal to 1. Note that, for the sake of simplicity, we assumed a diagonal corvariance matrix Λ and that *m*(**x**) was a linear combination of the fitness components 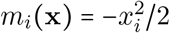. More general covariance matrices Λ and quadratic forms *m*(**x**) could have been considered as well using the transformations of the phenotype space presented in [29].

In the one-dimensional case, an equation of the form (4) with a general nonlocal reaction term of the form 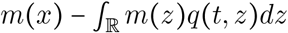 has been studied in [3]. Under the assumption that *m*(*x*) tends to -∞as *x* tends to ±∞, the authors have established a formula for *q* involving the eigenelements of the operator 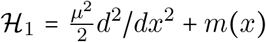. The formula can be made more explicit for our choice of *m*, *m*(*x*) = −*x*^2^/2 [2]. However, the method used in [2], which consists in reducing the equation (4) to the heat equation thanks to changes of variables based on Avron-Herbst formula and generalized Lens transform, cannot be directly applied in our *n*–dimensional anisotropic framework. Recently, another method based on constrained Hamilton-Jacobi equations has been developed to study the evolution of phenotypically structured populations, with integral or differential mutation operators (*e.g*., [6, 10, 15, 25, 35]). This method assumes a small mutation parameter *μ*^2^/2 (the terms 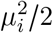 in (4)) of order *ε* ≪ 1, and is based on a scaling *t* → *t*/*ε*. Thus, it typically describes asymptotic evolutionary dynamics, at large times and in a ‘small mutation’ regime. Note however that, as *μ*^2^ = *U*λ, with *U* the mutation rate and λ the mutational variance (at each trait), it encompasses the cases where the mutation rate is not small, provided that λ ≪ 1/*U*. This method should apply with anisotropic mutation operators, as in (4). However, to the best of our knowledge, it cannot lead to explicit transient trajectories of adaptation, which is the real objective of our work. We use here a completely different approach.

In order to understand the dynamics of adaptation, an important quantity is of course the *fitness distribution p*(*t*, *m*), such that *p*(*t*, *m*)*dm* is the pushforward measure of the measure *q*(*t*, **x**)*d***x** by the map **x** ↦ −‖**x**‖^2^/2, and the *mean fitness* 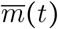. In the isotropic case [18], the authors directly focused on the equation satisfied by *p*(*t*, *m*) and not by *q*(*t*, **x**). Whereas (4) is quadratic into the variables *x*_1_,…, *x_n_*, the equation satisfied by *p*(*t*, *m*) is linear with respect to *m*, which makes possible the derivation of some PDEs satisfied by generating functions of *p*(*t*, *m*). Here, due to the anisotropy, the dynamics of *p*(*t*, *m*) is not summarized by a single one-dimensional PDE. Instead, we define the fitness components **m** = (*m*_1_,…, *m_n_*) and the *joint distribution of the fitness components* **p**(*t*, **m**) such that **p**(*t*, **m**)*d***m** is the pushforward measure of the measure *q*(*t*, **x**)*d***x** by the map 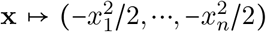 (see Proposition 2.3 for more details). As fitness is the sum of the fitness components, the distributions **p** and *p* are linked by:

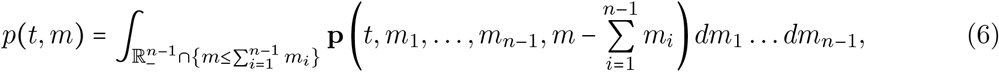

where ℝ_−_ = (−∞, 0].

Thus the mean fitness 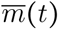 can be easily connected with these distributions:

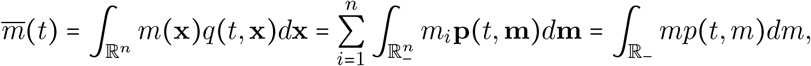

see also (15) below.

This paper is organized as follows. The results are presented in Section 2, and their proofs are presented in Section 4. More precisely, Section 2.1 is dedicated to the analysis of the time-dependent problem (4). We begin with the existence and uniqueness of the solution of the Cauchy problem. We then give an explicit formula for *q*(*t*, **x**) in the particular case of a Gaussian initial distribution of phenotypes *q*_0_(**x**). Then, we derive a nonlocal degenerate parabolic equation satisfied by **p**(*t*, **m**), and the equation satisfied by its cumulant generating function (CGF). Solving the equation satisfied by the CGF, we derive an explicit formula for 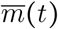. Then, in Section 2.2, we study the long time behavior and the stationary states of (4). We shall see that the distribution of the fitness components **p**(*t*, **m**) converges towards a distribution **p**_∞_(**m**) as *t* → +∞, and we give an explicit formula for **p**_∞_(**m**). Lastly, in Section 2.3 we show that including anisotropy in the models may help to understand experimental trajectories of fitnesses, such as those obtained in the famous experiment of Richard Lenski [23, 24, 45].

## 2 Main results

### 2.1 The time-dependent problem

#### Solution of the Cauchy problem associated with equation (4) for *q*(*t*, **x**)

We first show that the Cauchy problem admits a unique solution. We need the following assumption on the initial distribution *q*_0_:

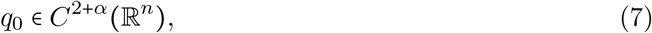

for some *α* ∈ (0,1), that is, 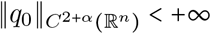. Additionally, we assume that:

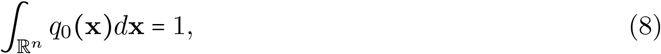

and there exists a function *g*: ℝ_+_ → ℝ_+_ (with ℝ_+_ = [0, +∞)) such that:
*g* is non-increasing, 0 ≤ *q*_0_ ≤ *g*(‖·‖) in ℝ^*n*^, **x** → *m*(**x**)*g*(‖**x**‖) is bounded in ℝ^*n*^,

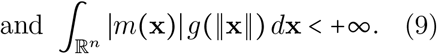

*These assumptions are made throughout the paper, and are therefore not repeated in the statements of the results below*.

We can now state an existence and uniqueness result for the distribution of phenotypes.

##### Theorem 2.1.

*There exists a unique nonnegative solution q* ∈ *C*^1,2^(ℝ_+_ × ℝ^*n*^) *of*:

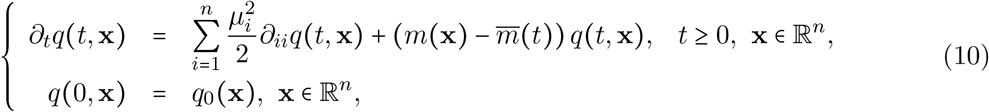

*such that q* ∈ *L*^∞^((0, *T*) × ℝ^*n*^) *for all T* > 0, *and the function*:

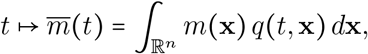

*is real-valued and continuous in* ℝ_+_. *Moreover, we have:*

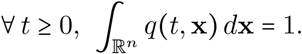

The next result gives an explicit solution of (10) in the particular case where the phenotypes are initially Gaussian-distributed.

##### Corollary 2.2.

Assume that the initial distribution of phenotype frequencies is Gaussian, that is,

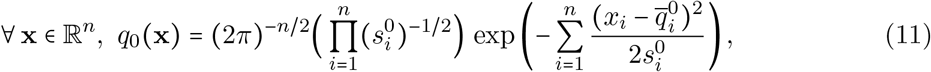

for some parameters 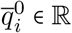 and 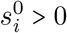. Then the solution *q*(*t*, **x**) of the Cauchy problem (10) is Gaussian at all time:

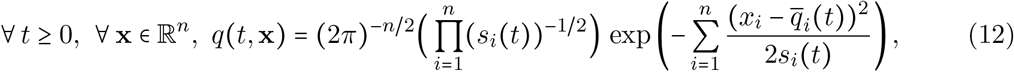

with:

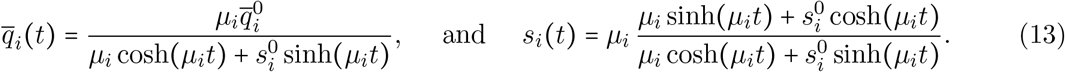

Moreover, we have:

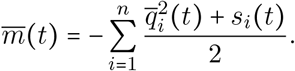

We also note, in Corollary 2.2, that the distribution *q*(*t*, **x**) converges to a Gaussian distribution with mean 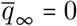 and variances *s*_∞,*i*_ = *μ_i_*, as *t* → +∞.

The determination of an explicit formula for *q*(*t*, **x**) becomes more involved when the initial distribution *q*_0_ is not a Gaussian. In this case, we study the equation satisfied by the distribution of the fitness components **p**(*t*, **m**) (see Introduction) and, as a by-product, we derive an explicit formula for 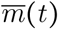.

#### A degenerate parabolic PDE satisfied by p(*t*, m)

Our objective here is to derive an equation for **p**(*t*, **m**) that only involves linear dependencies with respect to the coefficients *m_i_*, and holds for general initial phenotype distributions *q*_0_, which may not be Gaussian.

First, we express the distribution of the fitness components **p**(*t*, **m**) in terms of the distribution of phenotypes *q*(*t*, **x**) given in Theorem 2.1.

##### Proposition 2.3.

*For all t* ≥ 0 *and* 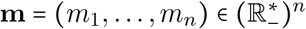, *there holds:*

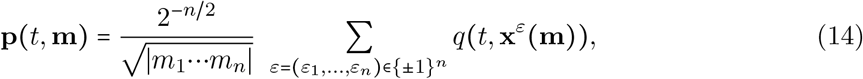

*with* 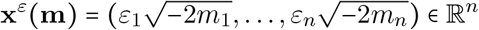. *Furthermore, we have:*

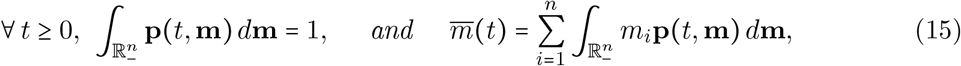

*where all above integrals are convergent*.

It also turns out that the expression (14) becomes simpler when *q* satisfies some symmetry properties. In the sequel, for a given function *f* ∈ *C*(ℝ^*n*^), we define its #-symmetrization *f*^#^ ∈ *C*(ℝ^*n*^) by:

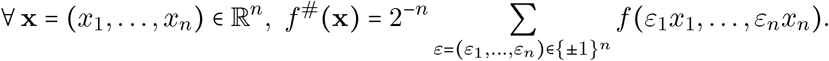

From the symmetries inherent to (4), it is easy to check that, if *q* is the solution of (4) with initial condition *q*_0_ satisfying the conditions of Theorem 2.1, then *q*^#^ is the solution of (4) with initial condition 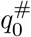, and *q*(*t*, ·) and *q*^#^(*t*, ·) have the same mean fitness 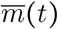 at every time *t* ≥ 0. Furthermore, using the expression (14), we observe that **p**(*t*, **m**) can be described in terms of the #-symmetrization of *q*(*t*, ·;):

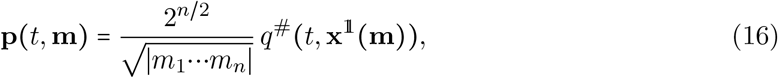

for every *t* ≥ 0 and 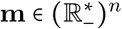, with:

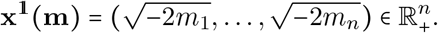

This function **p**(*t*, **m**) satisfies a degenerate parabolic equation:

##### Theorem 2.4.

*The distribution function of the fitness components* **p** *is a classical* 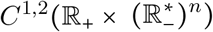 *solution of:*

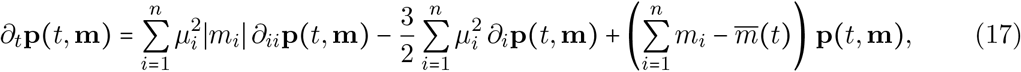

*for t* ≥ 0 *and* 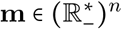, *with initial condition:*

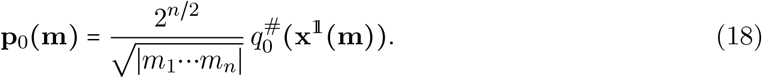

The equation (17) is degenerate in the sense that the operator in the right-hand side is not uniformly elliptic, as at least one of the coefficients 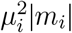 in front of the second order differential terms vanishes at the boundary of 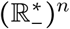.

In the isotropic case (*μ_i_* = *μ* > 0 for all *i*), it is also possible to derive a scalar equation for the distribution of fitness *p*(*t*, *m*), defined by (6).

##### Theorem 2.5.

(*Isotropic case*) *If μ_i_* = *μ for all* 1 ≤ *i* ≤ *n*, *then the fitness distribution p*(*t*, *m*) *is a classical* 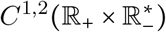 *solution of:*

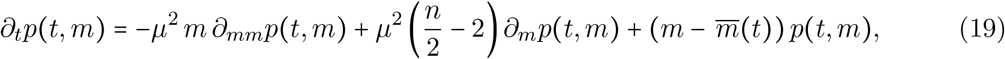

*for t* ≥ 0 *and m* < 0, *with initial condition*,

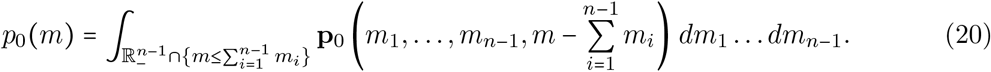

As expected, the equations (17) and (19) only involve linear combinations of the coefficients *m_i_*. This allows us to derive simpler equations satisfied by the generating functions of **p**(*t*, **m**) and *p*(*t*, *m*).

#### Generating functions

We define the *moment generating functions* (MGFs) *M*_**p**_ and *M_p_* of **p** and *p* and their logarithms – the *cumulant generating functions* (CGFs) – *C*_**p**_ and *C_p_* by:

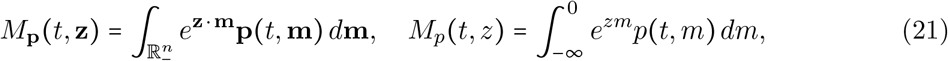

and:

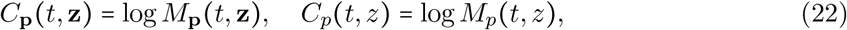

for *t* ≥ 0, 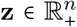 and *z* ≥ 0. The integrals are well defined because the components *m_i_* are all negative. Furthermore, it follows from (15) and the nonnegativity of **p** that, for each *t* ≥ 0, the functions *M*_**p**_(*t*, ·) and *C*_p_(*t*, ·) are of class 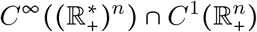, while the functions *M_p_*(*t*, ·) and *C_p_*(*t*, ·) are of class 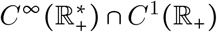.

The following result gives the equation satisfied by *C*_**p**_.

##### Theorem 2.6.

*The cumulant generating function C*_p_ *of* **p** *is of class* 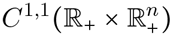^1^ *and it solves:*

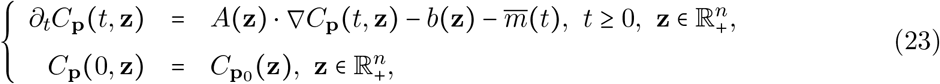

*where:*

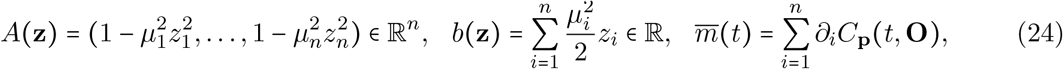

*and* ∇*C*_**p**_(*t*, **z**) *denotes the gradient of C*_**p**_ *with respect to the variable* **z**.

As a corollary of this theorem, we derive an equation satisfied by *C_p_* in the isotropic case.

##### Corollary 2.7.

(Isotropic Case) If *μ_i_* = *μ* for all 1 ≤ *i* ≤ *n*, then the cumulant generating function *C_p_* of the fitness distribution *p* is a *C*^1,1^(ℝ_+_ × ℝ_+_) solution of:

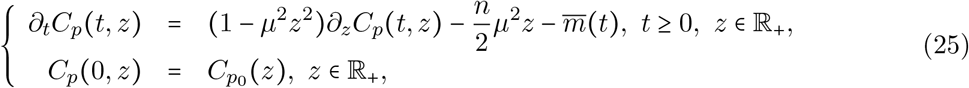

where *p*_0_ is the initial fitness distribution given in (20), and 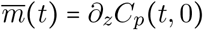.

Note that the equation (25) is directly obtained as a by-product of (23), without using the equation (19) satisfied by *p*(*t*, *m*). Each of these two equations (23) and (25) has a unique solution which can be computed explicitly, leading to a formula for 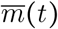. We refer to Section 2.3 for an application of this result.

In the Introduction, we mentioned that the coefficients 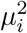 can be interpreted as the product between the mutation rate *U* and the variance λ_*i*_ at the *i*-th trait. In the isotropic case, 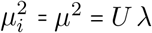. Thus we have retrieved the equation mentioned in [32, Eq. (E5) in Appendix E].

The last two results of this section provide some explicit expressions of *C*_**p**_(*t*, **z**) and *C_p_*(*t*, *z*) when **z** and *z* are close enough to **O** and 0.

##### Proposition 2.8.

*The cumulant generating function C*_**p**_ *of* **p** *is given by, for all t* ≥ 0 *and* **z** = (*z*_1_,…,*z**n***) ∈ [0,1/*μ*_1_) ×⋯× [0,1/*μ_n_*),

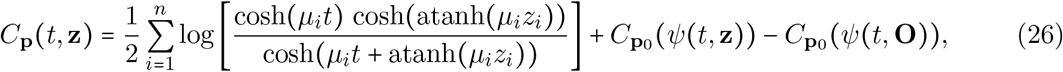

*with:*

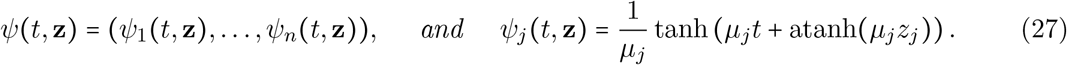

*Moreover, for all t* ≥ 0, *we have:*

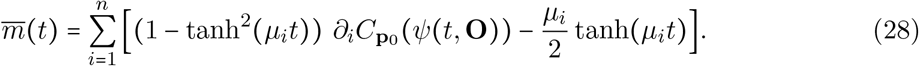

##### Corollary 2.9.

(Isotropic case) If *μ_i_* = *μ* for all 1 ≤*i* ≤ *n*, then the cumulant generating function *C_p_* of *p* is given by:

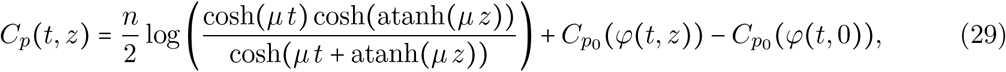

for *t* ≥ 0 and 0 ≤ *z* < 1/*μ*, with *φ*(*t*, *z*) = (1/*μ*) tanh (*μt* + atanh(*μz*)). Moreover, we have:

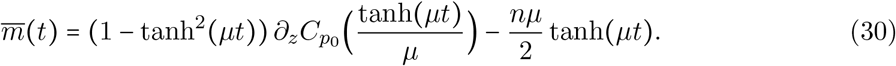

### 2.2 Long time behavior and stationary states

We are here interested in the long time behavior of the solutions of (4) and (17). We begin with a result on the convergence of the solution of (17) at *t* → +∞.

#### Theorem 2.10.

*Let* **p** *and* 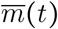 *be as in the previous section. Then:*

i. **p**(*t*, ·) *weakly converges in* 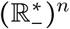 *to* **p**_∞_ *as t* → +∞, *where:*

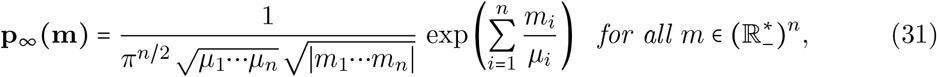

*in the sense that* 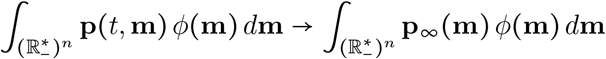 *as t* → +∞ *for every test function* 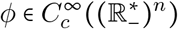;
ii. 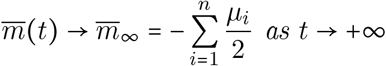 *and* 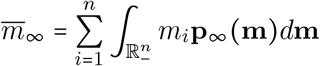;
iii. *the function* **p**_∞_ *is a classical* 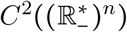 *solution of:*

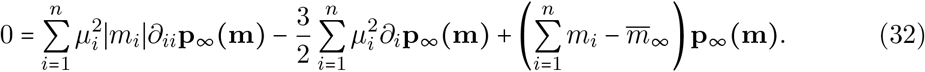 In the isotropic case, we retrieve the result of [32] in the WSSM case (Weak Selection and Strong Mutation), which says that the fitnesses are asymptotically distributed according to the symetrized Gamma distribution −Γ(*n*/2, *μ*), with 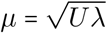:

#### Corollary 2.11.

(Isotropic case) If *μ_i_* = *μ* for all 1 ≤ *i* ≤ *n*, then *p*(*t*, ·) weakly converges in 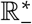 to *p*_∞_ as *t* → +∞, where:

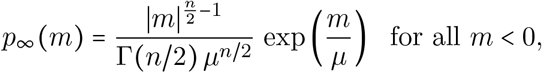

and 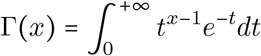 is the standard Gamma function.

Thanks to the previous two results, we get the asymptotic behavior of the phenotype distribution in the symmetric case.

#### Corollary 2.12.

If *q*_0_ is #-symmetric in the sense that 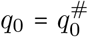, then *q*(*t*, ·) = *q*^#^(*t*, ·) weakly converges to *q*_∞_ as *t* → +∞ on ℝ_*n*_ where, for all *x* ∈ ℝ^*n*^,

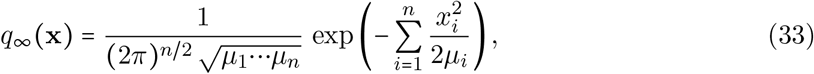

and 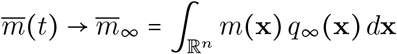 as *t* → +∞.

We note that *q*_∞_ is a classical positive stationary state of (4), *i.e*., it satisfies the following equation:

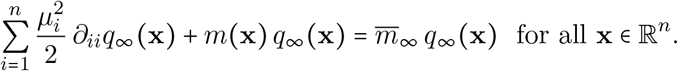

Thus, the distribution *q*(*t*, **x**) and the mean fitness 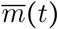 converge to the principal eigenfunction (resp. eigenvalue) of the operator 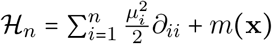.

In the 1D case (*x* ∈ ℝ), the results in [3] imply that *q*(*t*, *x*) can be written in terms of the eigenelements (λ_*k*_, *ϕ_k_*) of the operator 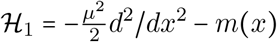. Namely,

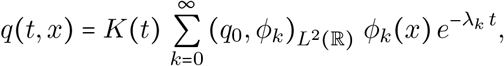

where *K*(*t*) is such that *q*(*t*, ·) sums to 1, and (·,·)_*L*^2^(ℝ)_ is the standard scalar product in *L*^2^(ℝ). As a corollary, the authors of [3] obtained the convergence of the distribution *q*(*t*, *x*) and of the mean fitness 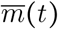 to the principal eigenfunction (resp. eigenvalue) of 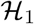, namely *ϕ*_0_(resp. λ_0_). Thus, in the 1D case, their result is stronger than that of Corollary 2.12, as the convergence of *q*(*t*, *x*) occurs in *L^p^*(ℝ^*n*^), for all 1 ≤ *p* ≤ +∞, and does not require *q*_0_ to be #-symmetric. Based on the spectral properties of the operator 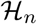 [20], we conjecture that their results can be extended to the anisotropic multidimensional framework considered here, meaning that the convergence result in Corollary 2.12 remains true when the initial condition *q*_0_ is not #-symmetric.

### 2.3 Effect of anisotropy: numerical computations and connection with *Escherichia coli* long-term evolution experiment

The objective of this section is to illustrate the importance of taking anisotropy into account when modelling adaptation trajectories. Isotropic models [32] lead to regularly saturating trajectories of 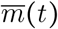 with a plateau, *i.e*. a single ‘epoch’. Here, we show that, in the presence of anisotropy, the trajectory of 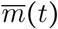 can exhibit several plateaus before reaching a stable level close to 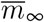. Thus, the dynamics of adaptation can show several evolutionary ‘epochs’, as those observed in the *E. coli* long-term evolution experiment [19], corresponding to different time-scales at which adaptation occurs.

For the sake of simplicity of the computations, and although the existence and uniqueness results of Section 2.1 were only obtained with continuous initial distributions of phenotypes, we assume a Dirac initial distribution of the phenotypes. Namely, we assume that *q*_0_ = *δ*_x_0__ with **x**_0_ = (*x*_0,1_,…,*x*_0,*n*_) ∈ ℝ^*n*^. This corresponds to an initially clonal population. This hypothesis implies that the initial distribution of the fitness components and the initial CGF are respectively given by:

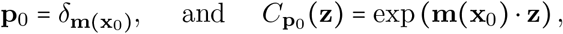

with 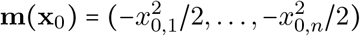. In this case, the expression (28) in Proposition 2.8 simplifies to:

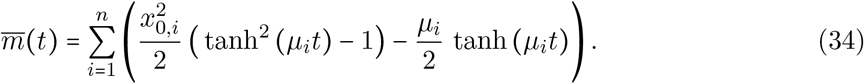

#### Trajectory of adaptation in the presence of anisotropy: an illustrative example

We take *n* = 3 and *μ*_1_ > *μ*_2_ > *μ*_3_. The corresponding trajectory of 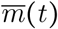 is depicted in Figure 1. After a brief initial decay which was already observed in the isotropic case [32], 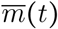 rapidly increases and reaches a first plateau (of value close to 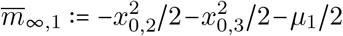). Then, 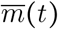 rapidly increases again to reach a second plateau (of value close to 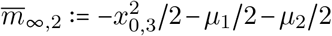). Finally, 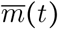 increases again and stabilises around 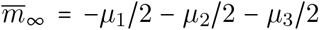. Interestingly, although the ultimate value 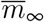 does not depend on the initial phenotype, the intermediate plateaus depend on **x**_0_. Their values approximately correspond to the fitnesses associated with the successive projections of **x**_0_ on the hyperplanes {*x*_1_ = 0} and {*x*_1_ = *x*_2_ = 0} minus the mutation load (we recall that the optimal phenotype was fixed at **x** = (0,…, 0), see Introduction).

**Figure 1 –.**
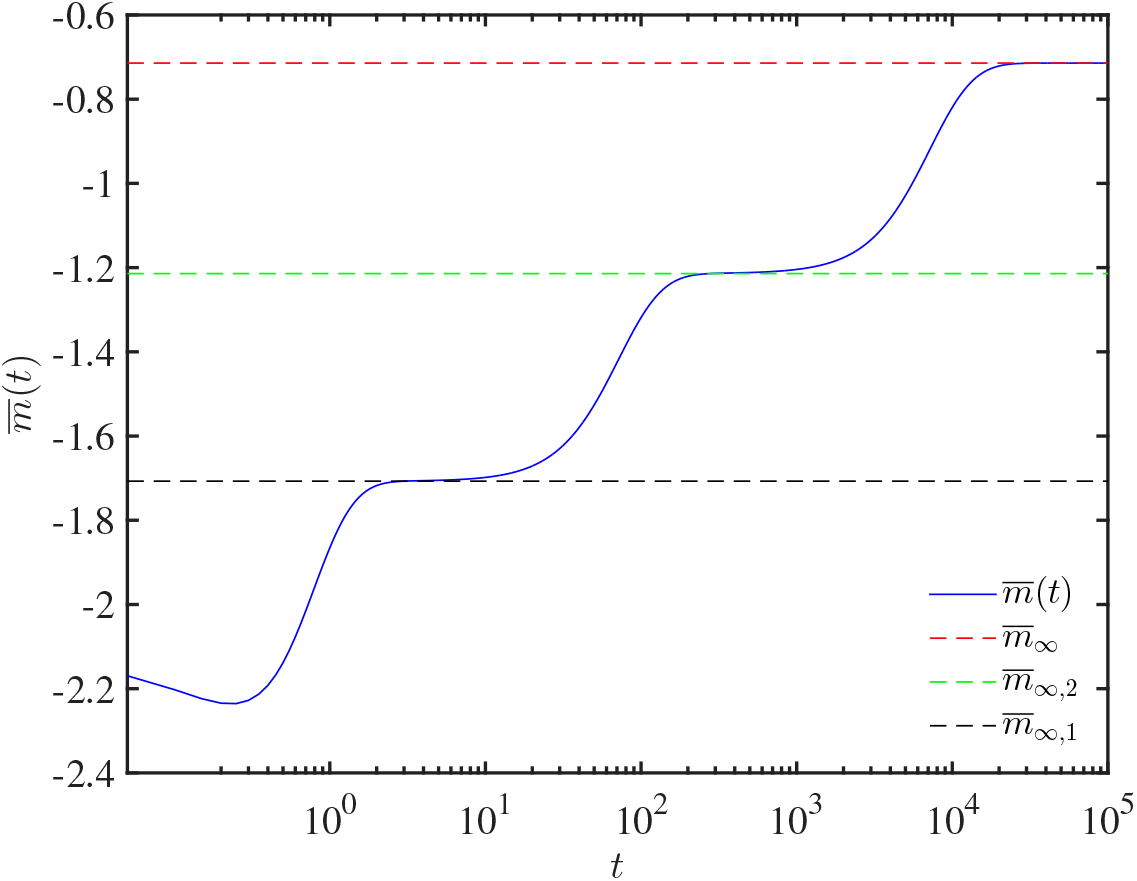
Trajectory of adaptation in the presence of anisotropy, with *n* = 3. The function 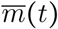 is given by formula (34), and the approximations of the values of the intermediate plateaus are 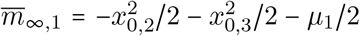 and 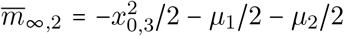. The mutational parameters are 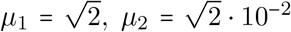 and 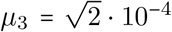. The other parameter values are **x**_0_ = (3/2,1,1).

More generally speaking, we prove in Section 4.5 that, for *n* ≥ 2 and *μ*_1_ > … > *μ_n_*, the trajectory exhibits (*n* − 1) plateaus (before the final one of value 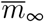), of respective values:

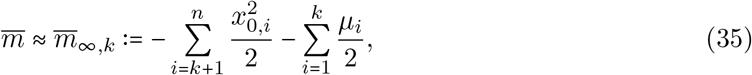

for *k* = 1,…, (*n* − 1). More precisely, we obtain the following proposition.

##### Proposition 2.13.

*Let* 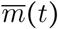 *and* 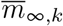 *be defined by* (34) *and* (35), *respectively. Given* **x**_0_ = (*x*_0,1_,…, *x*_0,*n*_) ∈ ℝ^*n*^, *T* > 0, *ε* > 0 *and μ*_1_ > 0, *there exist μ*_2_,…,*μ_n_* > 0 *such that, for each k* ∈ 〚 1, *n* − 1‛, *the set:*

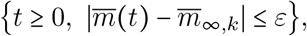

*contains an interval of length at least equal to T*.

Note that the values 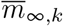 may not be ordered by increasing order, depending on the parameter values, possibly leading to nonmonotone trajectories of 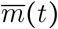. The plateaus are more visible when the *μ_i_*’s have different orders of magnitude. More precisely, we show in Section 4.5 that, given *T* > 0, 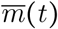 remains around each plateau of value 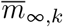 at least during a time interval of duration *T*, for a good choice of the parameters *μ_i_*.

##### Remark 2.14.

*The proof of Proposition* 2.13 *does not lead to explicit expressions for the intervals at which the plateaus occur. In the particular case n* = 2, *we can obtain a more precise characterization of these intervals, see Section 4.5*. *In particular, taking μ*_1_ = 1, *μ*_2_ = 10^−k^ *for some k* ≥ 1 *and x*_0,1_ = *x*_0,2_ = 1, *we show that:*

i. 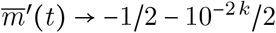 *as t* → 0;
ii. *at t* = *t*_0_ = ln(3), 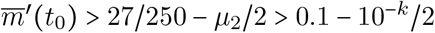;
iii. *for all t in the interval I*: = (ln (2 · 10^*k*^), 10^*k*/2^), 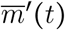 *is of order* 10^−3*k*/2^;
iv. *at* 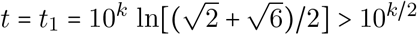, 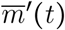 *is of order* 10^−*k*^.

*At larger times, we already know that* 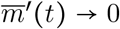. *Finally, this shows that* 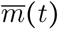 *remains stable within the interval I, corresponding to a part of the plateau, relatively to the times t*_0_ *(before the interval I) and t*_1_ *(after the interval I)*.

#### Long term evolution experiment with *Escherichia coli*

The long term evolution experiment (LTEE) has been carried by Lenski and his collaborators since 1988 [23]. Twelve populations of *E. coli* have been founded from a single common ancestor, and are still evolving after more than 70,000 generations. The fitness evolved rapidly during the first 2,000 generations, and then remained nearly static between generations 5,000 and 10, 000 [24], which would at least phenomenologically advocate for the existence of a phenotype optimum. However, more recent data (after generation 10,000) indicate that the mean fitness seems to increase without bounds [45]. Our goal here is not to propose a new explanation of the LTEE data, but simply to check whether the anisotropic model (4) leads to a better fit than an isotropic model.

The interpretation of the fitness data from the LTEE is quite subtle (see the comments in Richard Lenski’s Experimental Evolution website http://myxo.css.msu.edu/ecoli/srvsrf.html). For the sake of simplicity, as our objective is to check if the trajectories given by (34) can be qualitatively consistent with the fitness data from the LTEE, we make the following simplifying assumptions: (1) a time unit *t* in our model corresponds to 1 cycle of the experiment (1 day), corresponding to ≈6.64 generations [23]; (2) the link between 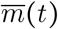 and the mean fitness values given by the LTEE data, which are of the form *w*(gen) (gen is the generation number and *w* the Darwinian mean fitness, which is related to the Malthusian fitness through an exponential) is made by assuming that 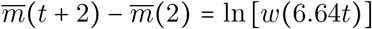. Thus, we assume (arbitrarily) that 2 cycles were necessary to build the founding colony from the single ancestor (hence the term *t*+2). Additionally, in the data, the fitness is a relative fitness against the ancestor, which implies that *w*(0) = 1; this is why the quantity 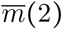 was subtracted to 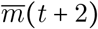. As mentioned above, the data are available for 12 populations. Here, we only use the data from one population (the population Ara-1, see [45]), for which measurements were obtained at 100-generation intervals during the first 2,000 generations, and then at 500-generation intervals.

We carried out a fit (using Matlab^®^ Curve Fitting Toolbox^®^, with a nonlinear least squares method) with the function 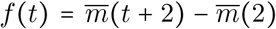. For the sake of simplicity, we assumed a two-dimensional phenotypic space (*n* =2). The only parameters to estimate are *μ*_1_, *μ*_2_and **x**_0_. We compared the result of this fit with a fit of the isotropic model (*μ*_1_=*μ*_2_). The results are depicted in Figure 2. A graphical analysis shows that the anisotropic model gives a far better fit. This is confirmed by the adjusted R^2^: 0.89 for the anisotropic model *versus* 0.57 for the isotropic model (R^2^ = 1 indicates that the fitted model explains all of the variability in the data). In the anisotropic model, the fitted mutational parameters have two different orders of magnitude: *μ*_1_ = 1.3 · 10^−2^and *μ*_2_=3.0 · 10^−4^. This leads to a first plateau until about 1,000 cycles (6,640 generations) followed by a second increase of the fitness. As expected, the isotropic model cannot explain this type of pattern.

**Figure 2 –.**
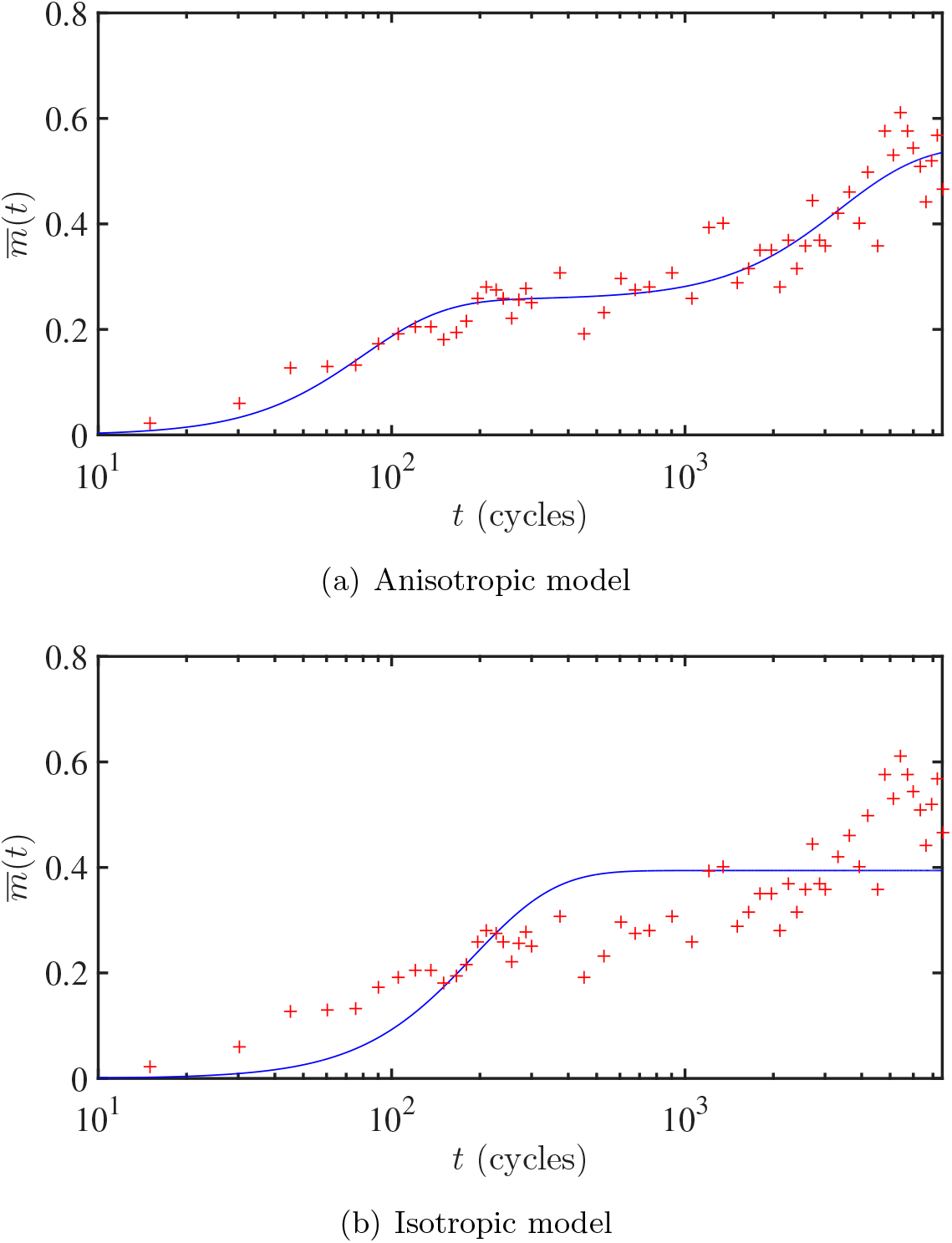
Trajectory of adaptation, anisotropic and isotropic model *versus* LTEE data. The function 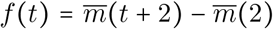, with 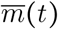 given by (34) was fitted to the LTEE data. The functions *f*(*t*) with the parameter values corresponding to the best fit are depicted in blue. The red crosses correspond to the LTEE data (population Ara-1, ln [*w*(6.64*t*)], with *w* the original data). The values leading to the best fit are (a, anisotropic model) **x**_0_ = (0.73,0.76), *μ*_1_ = 1.3 · 10^−2^ and *μ*_2_ = 3.0 · 10^−4^; (b, isotropic model) **x**_0_ = (0.30, 0.09) and *μ*_1_ = *μ*_2_ = 5.3 · 10^−3^.

## 3 Discussion

We considered a natural *n*-dimensional extension of the standard diffusive ‘replicator-mutator’ equation, to describe the dynamics of a phenotype distribution under anisotropic mutation and selection effects, and in the presence of a phenotype optimum. We proved that the corresponding Cauchy problem was well-posed (existence and uniqueness of the solution) and we proposed a new and general framework to the study of the quantitative behavior of the solution *q*(*t*, **x**). This framework enabled us to derive a formula for the mean fitness in the population at all time (equation (28)), an important quantity to describe the dynamics of adaptation.

The case of an initially Gaussian distribution *q*_0_ of the phenotypes is simpler, as the phenotype distribution *q*(*t*, **x**) remains Gaussian (though anisotropic) at all time. However, when *q*_0_ is not Gaussian, this result obviously breaks down. In this case, the method that we proposed consists in two steps: (1) to derive a degenerate nonlocal parabolic equation satisfied by the distribution **p**(*t*, **m**) of the fitness components, *i.e*., the joint distribution of the 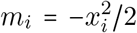. This equation is degenerate at 0, but has the advantage of being linear with respect to **m**; (2) to derive a nonlocal transport equation satisfied by the cumulant generating function of **p**(*t*, **m**). This last equation can be solved analytically, and its solution leads to explicit formulae for all the moments of **p**.

Conversely, the methods that are developed in this paper could be applied to solve more general degenerate parabolic PDEs. The idea would be firstly to transform the degenerate equation into a non-degenerate equation, through a change of function of the type (16) (rewriting *q*^#^ in terms of **p**) to get an existence result, and secondly to consider PDEs satisfied by moment generating functions in order to obtain uniqueness and more quantitative properties of the solution of the degenerate PDE.

A natural idea to solve (4) could be to consider directly the cumulant generating function associated with *q*:

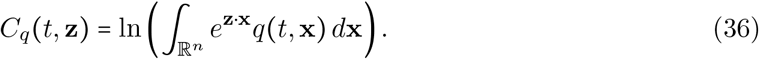

One would then have to see for which **z** this quantity makes sense, since now one integrates with respect to **x** ∈ ℝ_*n*_. Due to the nonlinear term *m*(**x**) in (4), the equation satisfied by *C_q_* would then be a nonlocal second-order viscous Hamilton-Jacobi type equation:

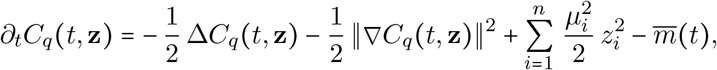

whereas equation (23) for *C*_**p**_ was a nonlocal first-order transport equation.

From an applied perspective, the results of Section 2.3 illustrate the importance of taking anisotropy into account, as it can open up further explanations of experimental data. In sharp contrast with the known results based on isotropic models, our results show that the trajectory of adaptation may exhibit (*n* − 1) plateaus before it reaches the final one. In particular, using our analytic formulae for the dynamics of 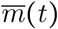, we obtained a very good fit of one of the most famous experimental dataset in experimental evolution, for which several evolutionary ‘epochs’ had already been observed [19]. This suggests that the FGM, in its anisotropic form, can generate a diversity of behaviours that may reconcile the various outcomes observed in long *versus* short term experimental evolution. However, whether this versatility may lead to overparameterization remains an open statistical question.

### Model extensions

The simple formalism of model (4) can easily be extended to more general situations. A first generalization corresponds to the case where the environment changes over time, which would be described by a moving optimum changing smoothly with some external environmental factor (*e.g*., a drug dose, a temperature, *etc*.). Such a moving optimum could be taken into account by setting:

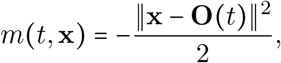

with **O**(*t*) ∈ ℝ^*n*^ the optimum at time *t*, instead of (5). In this case, we conjecture that Gaussian solutions of the form (12) still exist, suggesting that most our results could be extended to this situation.

Another generalization would be to explicitly consider birth-dependent mutation. Indeed, in many cases, mutations occur at birth. For example, in unicellulars undergoing a birth-death process, divisions without mutation occur at per capita rate *b*(1 − *u*) and divisions with mutation occur at rate *bu* where *u* is a per division mutation probability. If the birth rate varies across genotypes or over time, then so does the mutation rate, and we can no longer consider a constant per capita mutation rate *U*. First, let us note that this complication can be ignored in several biologically relevant situations:

- In multicellulars, evolving over a discrete time demographic process (*e.g*. annual plants), the process of birth and mutations are decorrelated as what determines the mutation rate is the number of divisions in the germline, which does not a priori correlate with fitness (fecundity or survival). Hence the model with constant *U* can be applied here, our model being a continuous time approximation to the exact discrete time model.
- In many microbial evolution experiments (like the LTEE presented in Section 2.3), cells undergo rounds of a pure birth process separated by regular dilutions by a constant factor. In this situation, if cells grow to a constant density, then remain ‘dormant’ before dilution (so-called stationary phase in microbiology) then each cycle corresponds to a constant number of divisions. With a faster birth rate, these divisions simply occur earlier on during the cycle, but are not more numerous. This process becomes approximately equivalent to a discrete time life cycle at the scale of growth dilution cycles. The mutation rate per cycle is then also constant (independent of the birth rate). Note that this heuristics is only approximate, as it ignores variation in the birth rate between genotypes, relative to its mean, among co-segregating lineages, over one growth cycle. However, that this heuristics applies to the LTEE may explain why models that ignore the coupling of birth and mutation rate can provide a good fit to data from the LTEE. More detailed individual-based stochastic models for the LTEE can be found in *e.g*. [5, 7].
- Under ‘viability selection’, selection only acts to reduce the death rate *d*, while the birth rate *b* shows limited evolution: mutation rates are then also roughly constant over time (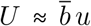, where 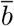 is the mean birth rate, stable over time). This happens if (i) the environmental challenge mainly affects the death rate and (ii) birth and death rates are uncorrelated so that evolution of *d* has little impact on *b*. In the context of phenotype-fitness landscape models, one situation where this applies is when fitness only depends on phenotype through the death rate, *m*(**x**) = *b* − *d*(**x**): the mutation rate is then *U* = *bu*, stable over time and across genotypes/phenotypes.

Overall there are several situations where one could ignore the connection between birth and mutation rates. Yet, in a more general context where the birth rate *b*(**x**) ≥ 0 does depend on phenotype **x**, then the mutation rate *per capita* per unit time also depends on **x**: *U*(**x**) = *ub*(**x**), with *u* the probability of mutation per birth event. The model (4) can then be extended, to take into account this effect:

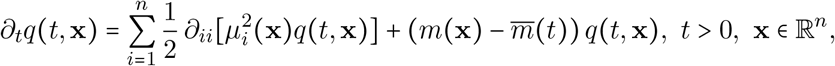

with 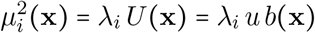, with λ_*i*_ the mutational variance at each trait, see Appendix A. However, the mathematical results of our paper cannot be straightforwardly extended to this case, even in the isotropic case. In particular, as soon as *b*(**x**) is not constant, this equation has no Gaussian solution. Still, characterization of the steady states should be possible as a principal eigenvalue problem.

## 4 Proofs

This section is devoted to the proofs of the results announced in Section 2. Section 4.1 is concerned with problem (10) satisfied by the distribution of phenotype frequencies *q*(*t*, **x**), while Section 4.2 deals with the fitness frequencies **p**(*t*, **m**) and the proofs of Proposition 2.3 and Theorem 2.4. In Section 4.3, we carry out the proofs of Theorem 2.6 and Proposition 2.8 on the cumulant generating functions, and their corollaries (Theorem 2.5 and Corollaries 2.7 and 2.9) in the isotropic case. Lastly, Sections 4.4 and 4.5 are concerned with the stationary states and the existence of plateaus for the mean fitness.

### 4.1 Proofs of Theorem 2.1 and Corollary 2.2 on the Cauchy problem (10)

Before considering the nonlocal problem (10), we begin with the local problem:

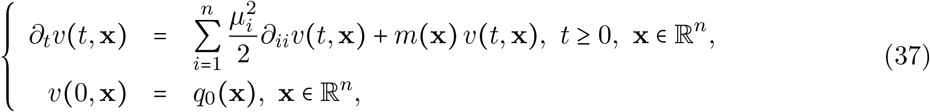

where *q*_0_ satisfies (7)–(9). As the fitness function:

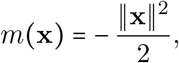

is unbounded, standard parabolic theory with bounded coefficients does not apply. However, some properties of (37) can be for instance be found in [4, 26, 36], which in particular lead to the following result.

#### Theorem 4.1.

*The problem* (37) *admits a unique bounded solution v* ∈ *C*^1,2^(ℝ_+_ × ℝ^*n*^). *Additionally, we have:*

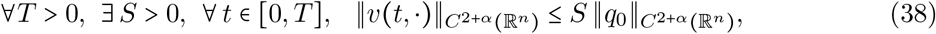

*and:*

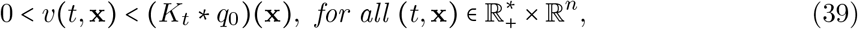

*with:*

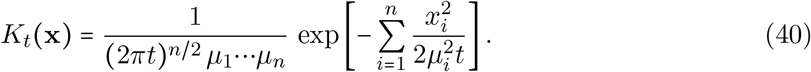

*Proof*. Let us fix a time *T* > 0. Theorem 2 in [26] implies that (37) admits a unique bounded solution *v* ∈ *C*^1,2^([0, *T*] × ℝ^*n*^) and this solution satisfies (38). Theorem III in [4] further implies that this solution is nonnegative. As *T* was chosen arbitrarily, these existence, uniqueness and nonnegativity results extend to *t* ∈ (0, +∞), with local boundedness in *t*.

Let us set *h*(*t*, **x**): = (*K_t_* * *q*_0_)(**x**) for *t* > 0, and *h*(0, **x**) = *q*_0_(**x**). The function *h* satisfies:

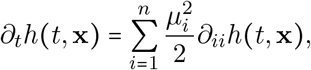

for all *t* > 0 and **x** ∈ ℝ^*n*^. Let *ψ*(*t*, **x**): = *v*(*t*, **x**) − *h*(*t*, **x**). We see that, for all *t* > 0 and **x** ∈ ℝ^*n*^,

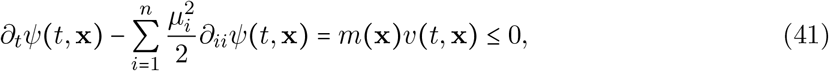

and *ψ*(0, **x**) = 0. By the Phragmen-Linderlöf principle [36, Theorem 10, Chapter 3], we get that *ψ* ≤ 0 in ℝ_+_ × ℝ^*n*^, *i.e*., *v*(*t*, **x**) ≤ *h*(*t*, **x**) = (*K_t_* * *q*_0_)(**x**). We therefore infer that 0 ≤ *v*(*t*, **x**) ≤ (*K_t_* * *q*_0_)(**x**) in (0, +∞) × ℝ^*n*^. Since *q*_0_ is bounded, this also implies that *v* is bounded in ℝ_+_ × ℝ^*n*^.

By the standard strong parabolic maximum principle, we conclude that the first inequality is strict, *i.e*., 0 < *v*(*t*, **x**), for all (*t*, **x**) ∈ (0, +∞) × ℝ^*n*^, since *v*(0, ·) = *q*_0_ is continuous, nonnegative and not identically equal to 0. Furthermore, since the inequality in (41) is then strict for all (*t*, **x**) ϵ (0, +∞) × (ℝ^*n*^ \ {**O**}), we get that *ψ*(*t*, **x**) < 0, *i.e*., *v*(*t*, **x**) < (*K_t_* * *q*_0_)(**x**), for all (*t*, **x**) ϵ (0, +∞) × ℝ^*n*^.

In order to connect (37) and (10), we need the following lemma.

#### Lemma 4.2.

The function:

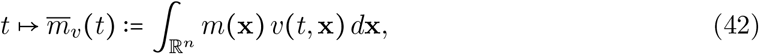

is real-valued and continuous in ℝ_+_ and, for every *t* ≥ 0, there holds:

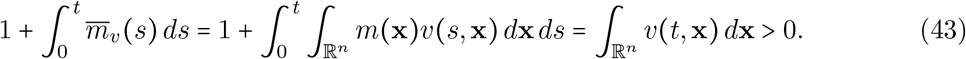

*Proof*. First of all, denote:

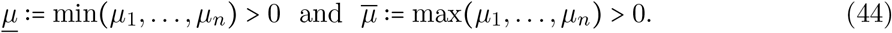

It follows from the assumptions on *q*_0_ that 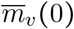 is a nonpositive real number. Consider now any *t* > 0 and let us check that 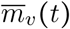 defined in (42) is a nonpositive real number. The function **x** → *m*(**x**) *v*(*t*, **x**) is nonpositive and continuous in ℝ^*n*^. Furthermore, it follows from Theorem 4.1 and the assumptions on *q*_0_ that:

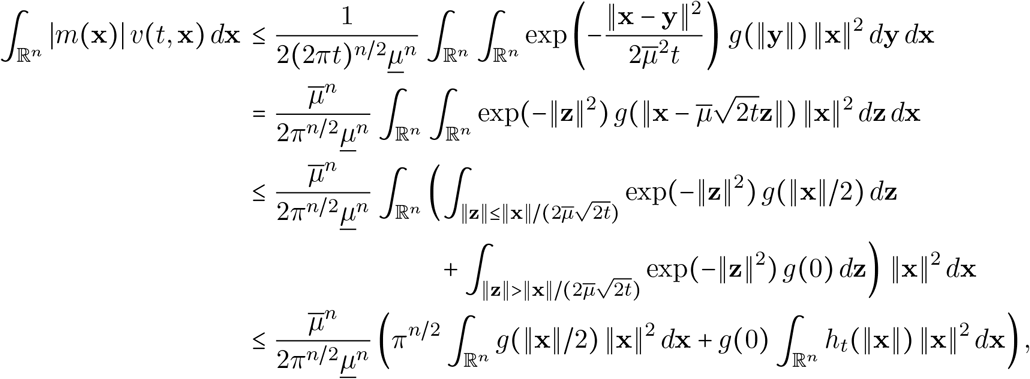

where:

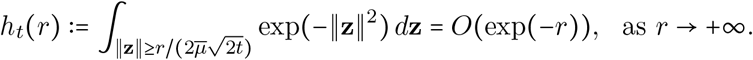

Since the integral 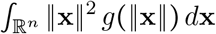 converges by assumption, one concludes that the integral 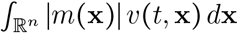 converges as well, hence 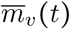 is a nonpositive real number. Furthermore, since the quantities *h_t_*(*r*) are non-decreasing with respect to *t* > 0, the same arguments together with Lebesgue’s dominated convergence theorem imply straightforwardly that the function 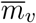 is continuous in ℝ_+_.

The convergence of the integral defining 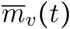 for every *t* ≥ 0, together with the nonnegativity and continuity of *v* and **x** → −*m*(**x**) = ‖**x**‖^2^/2, immediately implies that the integral 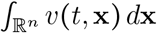 is a nonnegative real number for each *t* ≥ 0. Moreover, since *v*(*t*, **x**) > 0 for all 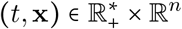, we infer that:

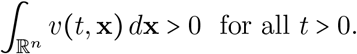

As in the previous paragraph, the function 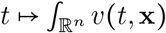 is also continuous in ℝ_+_.

Fix now an arbitrary *t* > 0. For *R* > 0, denote *B_R_* = {**x** ϵ ℝ^*n*^, ‖**x**‖ < *R*}, *ν* the outward unit normal on *∂B_R_* and *dσ*(**x**) the surface measure on *∂B_R_*. For every *ε* ϵ (0, *t*) and every *R* > 0, the integration of (37) over [*ε*, *t*] × *B_R_* yields:

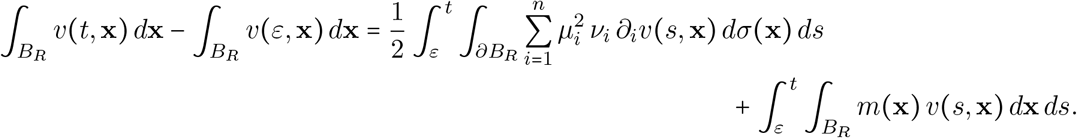

Since the first term of the right-hand side converges to 0 as *R* → +∞ from standard parabolic estimates (see [14]), one gets that:

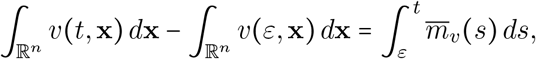

by passing to the limit *R* → +∞. The limit *ε* → 0^+^ then yields:

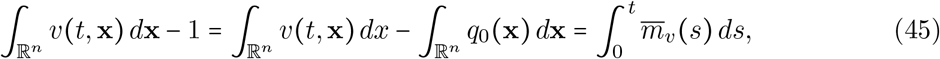

which gives the desired result (43) for *t* > 0. Formula (43) for *t* = 0 is trivial since *v*(0, ·) = *q*_0_ has unit mass. The proof of Lemma 4.2 is thereby complete.

#### Proof of Theorem 2.1.

Let *v* ≥ 0 be the unique classical solution of (37) given in Theorem 4.1. From Lemma 4.2, the function 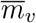 defined by (42) is continuous in ℝ_+_. Let us then set:

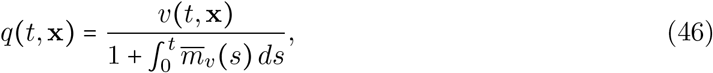

for every (*t*, **x**) ϵ ℝ_+_ × ℝ^*n*^. We recall that the denominator in the right-hand side of (46) is positive from Lemma 4.2. From Theorem 4.1 and Lemma 4.2, the function *q* is nonnegative and of class *C*^1,2^(ℝ_+_ × ℝ^*n*^). Furthermore, *q*(0, ·) = *v*(0, ·) = *q*_0_ in ℝ^*n*^ and it follows from Lemma 4.2 that:

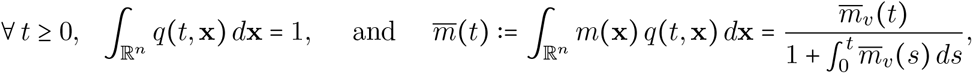

hence the function 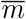 is real-valued, nonpositive and continuous in ℝ_+_. Lastly, it is immediate to see that *q* obeys (10). Furthermore, since *v* is bounded in (0, *T*) × ℝ^*n*^, the function *q* is bounded in (0, *T*) × ℝ^*n*^ for every *T* > 0.

To show the uniqueness, assume now that we have two nonnegative classical solutions *q*_1_ and *q*_2_ of (10) in *C*^1,2^(ℝ_+_ × ℝ^*n*^) ∩ *L*^∞^((0, *T*) × ℝ^*n*^) (for every *T* > 0), with the same initial condition *q*_0_ satisfying (7)–(9), and such that the functions:

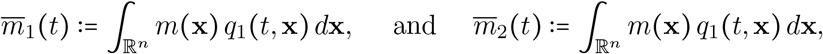

are real-valued and continuous in ℝ_+_. Define:

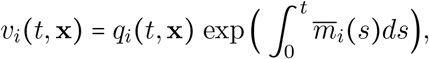

for *i* = 1,2, and (*t*, **x**) ϵ ℝ_+_ × ℝ^*n*^. The two functions *v*_1_ and *v*_2_ satisfy (37) and are bounded in (0, *T*) × ℝ^*n*^ for every *T* > 0. From Theorem 4.1, we get *v*_1_ ≡ *v*_2_ in ℝ_+_ × ℝ^*n*^. Furthermore, for all *i* = 1,2 and *t* ϵ ℝ_+_, there holds:

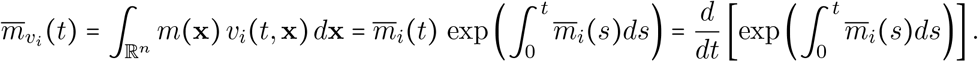

Hence, for all (*t*, **x**) ϵ ℝ_+_ × ℝ^*n*^, we get:

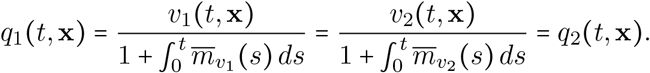

The proof of Theorem 2.1 is thereby complete.

#### Proof of Corollary 2.2.

It is a straightforward calculation to check that the function *q* defined by (12)–(13) is a classical bounded solution of (10) with initial condition given by (11). The conclusion then follows from the uniqueness part of Theorem 2.1.

### 4.2 A degenerate parabolic PDE satisfied by p(*t*, m)

#### Proof of Proposition 2.3.

We recall that the phenotypes are represented by *n* traits, and so by a vector in ℝ^*n*^ and that we have a constant optimum **O**, which is fixed to (0,…, 0) up to translation. We define, for each *ε* = (*ε*_1_,…,*ε_n_*) ϵ { ± 1}^*n*^, the subset:

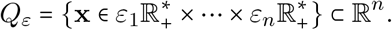

For any time *t* ≥ 0, we get from the law of total probability that:

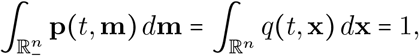

and that, for any 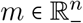,

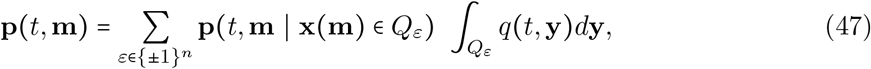

with **p**(*t*, **m**| **x**(**m**) ϵ *Q_ε_*) the conditional density of the fitness vector **m**, given that the associated phenotype **x**(**m**) is in *Q_ε_*. In the above formula (47), we also use the fact that 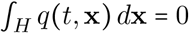 with:

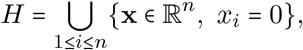

since *q*(*t*, ·) is continuous in ℝ^*n*^.

As the fitness function **x** ϵ *Q_ε_* ↦ **m**(**x**) = (*m*_1_(**x**),…, *m_n_*(**x**)) with 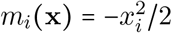 is one-to-one from *Q_ε_* to 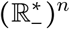, with inverse 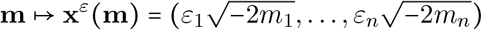, we infer that, for every *t* ≥ 0 and 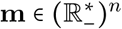,

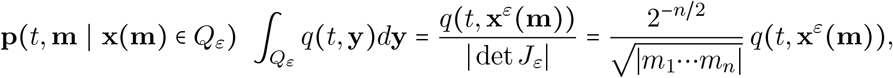

with 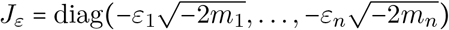. Finally, we get:

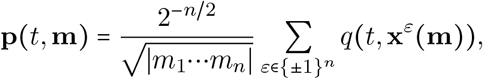

and:

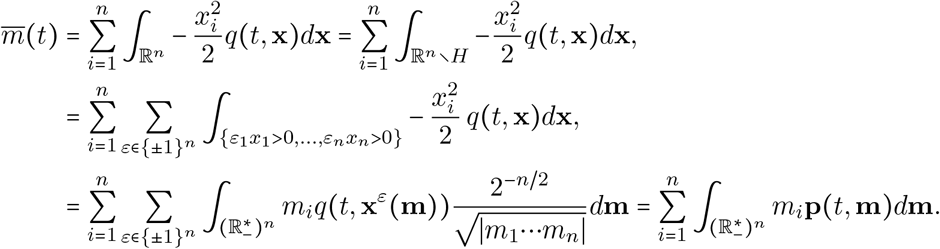

Notice that all integrals in the last sum converge since all integrands are nonpositive and the sum of these integrals is a real number. Observe lastly that 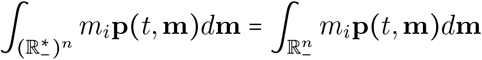, for every 1 ≤ *i* ≤ *n*, since **p**(*t*, ·) is an 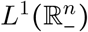 function. The proof of Proposition 2.3 is thereby complete.

#### Proof of Theorem 2.4.

Formula (16) implies that the function **p** is of class 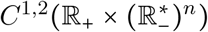 with initial condition **p**_0_ given by (18). Furthermore, it is straightforward to check that, for all *t* ≥ 0 and 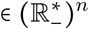:

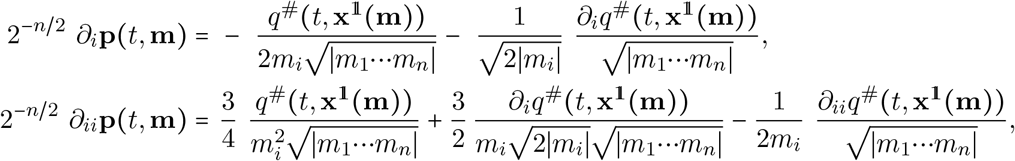

with 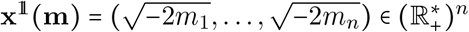. Hence, we have:

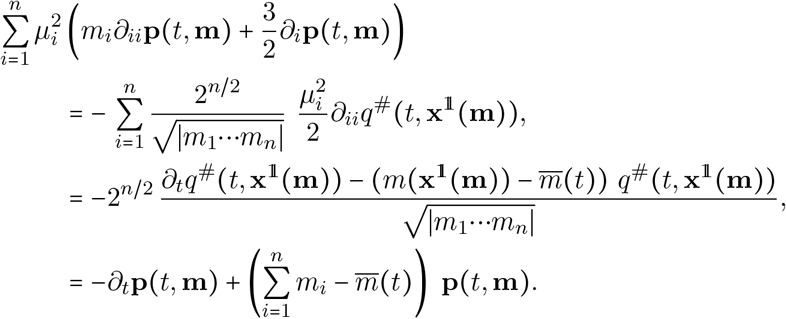

Theorem 2.4 is thereby proven.

#### Proof of Theorem 2.5.

Since by definition *p*(*t*, *m*) *dm* is the pushforward measure, at each time *t* ≥ 0, of the measure *q*(*t*, **x**) *d***x** by the map **x** ↦ −‖**x**‖^2^/2, it follows from the layer-cake formula that:

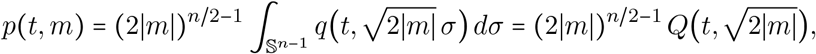

for all 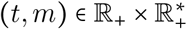, where 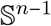 denotes the unit Euclidean sphere of ℝ^*n*^ and:

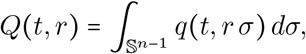

for (*t*, *r*) ϵ ℝ_+_ × ℝ_+_. Since *q* is of class *C*^1,2^(ℝ_+_ × ℝ^*n*^), it is easy to see that the function:

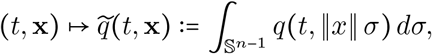

is of class *C*^1,2^(ℝ_+_ × ℝ^*n*^) too, hence the function *Q* is of class *C*^1,2^(ℝ_+_ × ℝ_+_) and *p* is of class 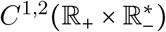. Furthermore, 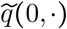 is of class *C*^2+*α*^(ℝ^*n*^) since *q*(0, ·) = *q*_0_ is of class *C*^2+*α*^(ℝ^*n*^).

Since *q* solves (10), which is invariant by rotation in the present isotropic case (*μ_i_* = *μ* for every 1 ≤ *i* ≤ *n*) and since 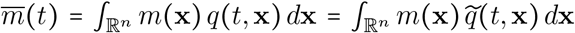, for every *t* ≥ 0, it follows that 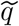 solves (10) as well, with initial condition 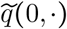. As a consequence, *Q* satisfies:

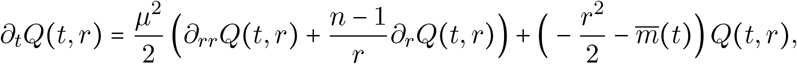

for all 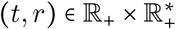. But since:

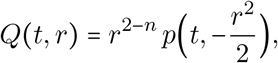

for all 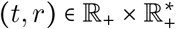, it is then straightforward to check that *p* solves (19) in 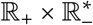. Lastly, the formula (20) is an immediate consequence of the definitions of *p* and **p**, and the proof of Theorem 2.5 is thereby complete.

### 4.3 Generating functions

#### Proof of Theorem 2.6.

Given *q*_0_ satisfying (7)–(9), we call *v* the unique bounded nonnegative *C*^1,2^(ℝ_+_ × ℝ^*n*^) solution of (37) defined in Theorem 4.1 with initial condition 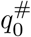. Notice that the function 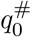 satisfies the same assumptions (7)–(9) as *q*_0_. By uniqueness and symmetry of (37) with respect to the change of variable *x_i_* into −*x_i_*, for any 1 ≤ *i* ≤ *n*, it follows that *v*(*t*, **x**) = *v*^#^(*t*, **x**) and, as in (46),

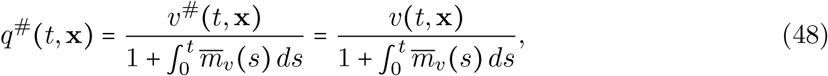

for all 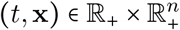.

As already noticed in Section 2.1, from (15) and the nonnegativity of **p**, the functions (*t*, **z**) ↦ *M*_**p**_(*t*, **z**) and (*t*, **z**) ↦ *C*_**p**_(*t*, **z**) given in (21)–(22) are well defined in 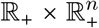. Let us start by proving the continuity of these functions *M*_**p**_ and *C*_p_ in 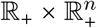. Owing to the relations (16) and (48), we see that:

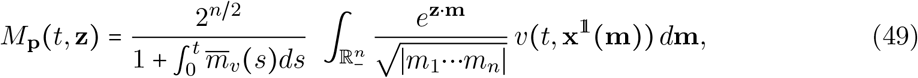

for all 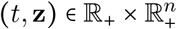, with 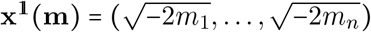. Hence, we have:

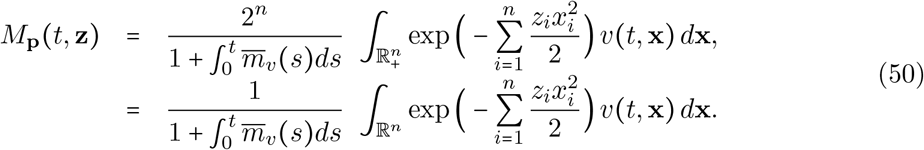

Notice that the function 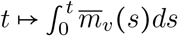 is continuous in ℝ_+_. Furthermore, the function 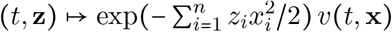 is also continuous in 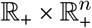, for every **x** ϵ ℝ^*n*^. Lastly, as in the proof of Lemma 4.2, it follows from (9), (39)–(40) and (44) that, for any 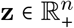, *T* > 0, *t* ϵ (0, *T*] and **x** ϵ ℝ^*n*^, there holds:

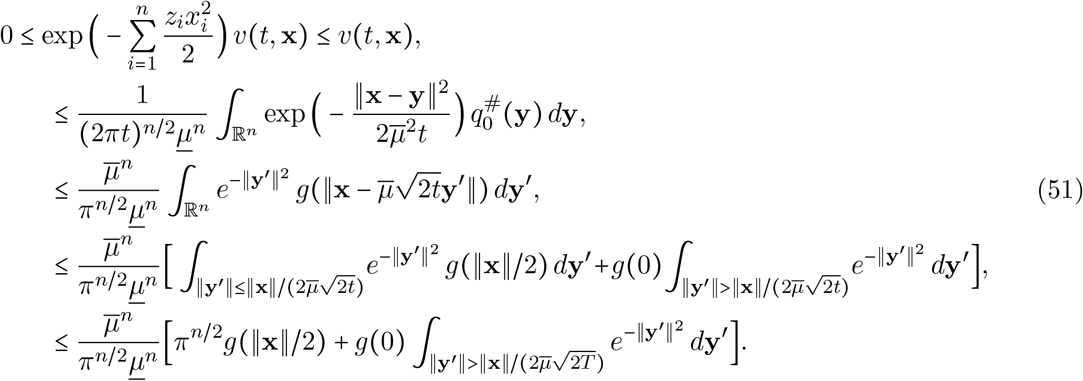

Call *h*(**x**) the quantity given in the right-hand side of the last inequality. Since by (9) the function *g*(‖·‖) is in *L*^∞^(ℝ^*n*^) ∩ *L*^1^(ℝ^*n*^), the function *h* belongs to *L*^1^(ℝ^*n*^), and is independent of 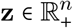 and *t* ϵ (0, *T*]. One then infers from (50) and Lebesgue’s dominated convergence theorem that the function *M*_**p**_ is continuous in 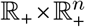. As *C*_**p**_ = log *M*_**p**_, the cumulant generating function *C*_**p**_ is also continuous in 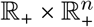.

Let us then check that *M*_**p**_ and *C*_**p**_ are 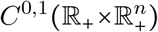, meaning that the functions 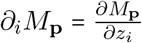 and 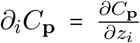 exist and are continuous in 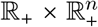, for every 1 ≤ *i* ≤ *n*. As a matter of fact, since **p**(*t*, ·) is a probability density function in 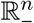 for any *t* ≥ 0 and since the integral 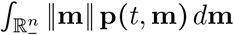 converges by formula (15) in Proposition 2.3, it easily follows from Lebesgue’s dominated convergence theorem that *∂_i_M*_**p**_(*t*, **z**) exists for all 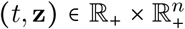 and 1 ≤ *i* ≤ *n*, with:

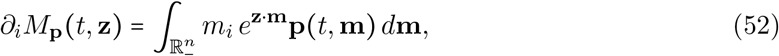

hence, as in (49)–(50),

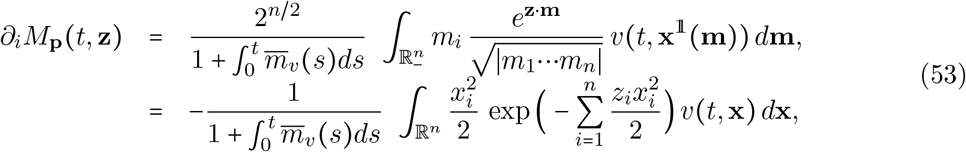

from (16) and (48). On the one hand, the function 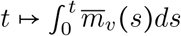 is continuous in ℝ_+_ and so is the function 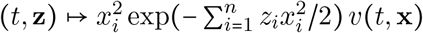 in 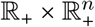, for every **x** ϵ ℝ^*n*^. On the other hand, as in the previous paragraph, it follows from (39)–(40) and (44) that, for any 1 ≤ *i* ≤ *n*, 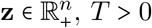, *t* ϵ (0, *T*] and **x** ϵ ℝ^*n*^, there holds:

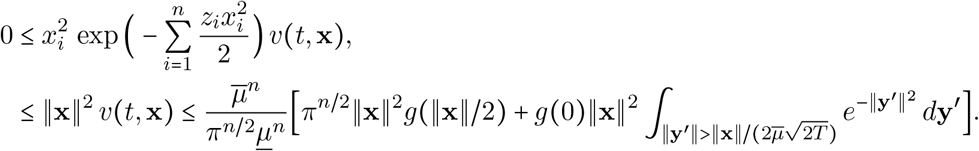

Call 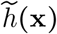 the quantity given in the right-hand side of the last inequality. Since by (9) the function **x** ↦ ‖**x**‖^2^ *g*(‖**x**‖) is in *L*^1^(ℝ^*n*^), the function 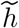 belongs to *L*^1^(ℝ^*n*^), and is independent of 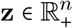 and *t* ϵ (0, *T*]. One then infers from (53) and Lebesgue’s dominated convergence theorem that the function *∂_i_M*_**p**_ is continuous in 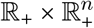, and so is the function *∂_i_C*_**p**_ = *∂_i_M*_**p**_/*M*_**p**_.

In this paragraph, we are interested in the differentiation of *M*_**p**_ with respect to *t*. By (37), we already know that:

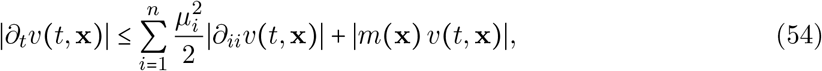

for all (*t*, **x**) ϵ ℝ_+_ × ℝ^*n*^. Fix *T* > 0 and let *S* > 0 be the constant given in (38). Thus, for all (*t*, **x**) ϵ [0, *T*] × ℝ^*n*^, there holds:

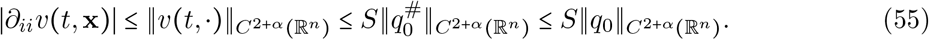

Let us now focus on the boundedness of the second term of the right-hand side of (54), that is, the boundedness of the function (*t*, **x**) → *m*(**x**)*v*(*t*, **x**) in [0, *T*] × ℝ^*n*^. Since this function is continuous in ℝ_+_ × ℝ^*n*^, let us show its boundedness in (0, *T*] × ℝ^*n*^. Thanks to (9) and (39)–(40), we get, as in (51), that:

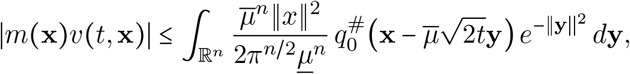

for all (*t*, **x**) ϵ (0, *T*] × ℝ^*n*^. Thus, we have:

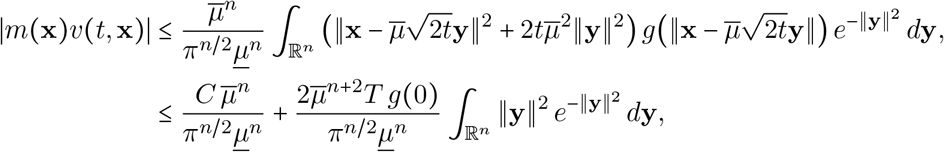

where the constant *C* is such that 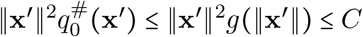, for all **x**′ ϵ ℝ^*n*^. Therefore, the function (*t*, **x**) ↦ *m*(**x**)*v*(*t*, **x**) is bounded in [0, *T*] × ℝ^*n*^ for any *T* > 0, and so is *∂_t_v* by (54)–(55). Together with (38), (48) and the continuity of 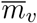 in ℝ_+_, it follows that the function *∂_t_q*^#^ is bounded in [0, *T*] × ℝ^*n*^, for every *T* > 0. Finally, (16) implies that for all 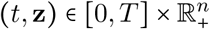 and 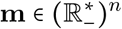,

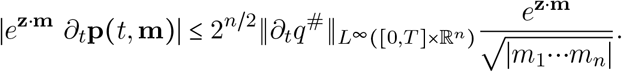

Since the integrals:

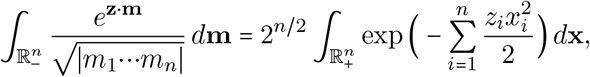

converge for all 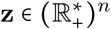, it then easily follows from the previous estimates and from Lebesgue’s dominated convergence theorem that the function *M*_**p**_ is differentiable with respect to *t* in 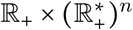, with:

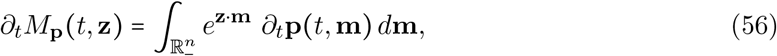

and that the function *∂_t_M*_**p**_ is itself continuous in 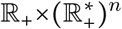. So is the function *∂*_*t*_*C*_**p**_ = *∂_t_M*_**p**_/*M*_**p**_. The continuity of the functions *∂_t_M*_**p**_ and *∂_t_C*_**p**_ in the closure 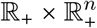 of 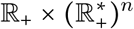 will be obtained as a consequence of the equations satisfied by these two functions, which shall be established below.

Let us then turn to find an equation satisfied by *M*_**p**_, in order to derive the equation (23) satisfied by *C*_**p**_. Fix 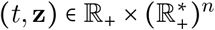. Thanks to (17), we have:

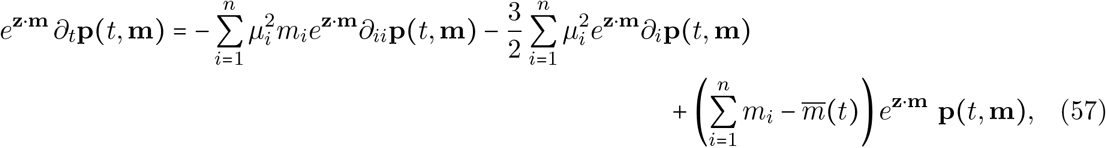

for all 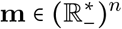.

We are now going to integrate (57) over 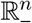. To do so, let us first focus on the first two terms of the right-hand side of (57). Fix an index *i* ϵ 〚 1,*n*〛 and consider the cubes:

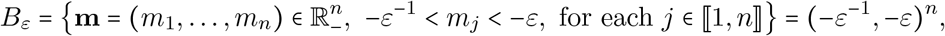

with 0 <ε < 1. Denote 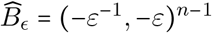 and:

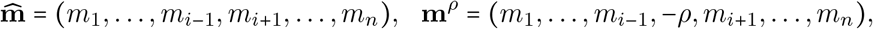

for *ρ* ϵℝ. By using Fubini’s theorem and integrating by parts with respect to the variable *m_i_*, one infers that:

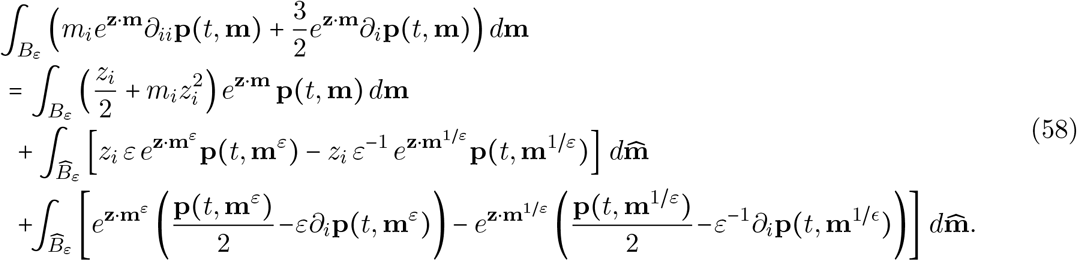

Let us pass to the limit as *ε* → 0 in the three integrals of the right-hand side of (58). Firstly, since **p**(*t*, ·) is nonnegative and the functions **m**↦**p**(*t*, **m**) and **m**↦*m_i_***p**(*t*, **m**) are in 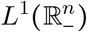, it follows from Lebesgue’s dominated convergence theorem together with (21) and (52) that:

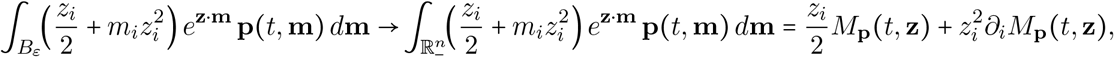

as *ε* → 0. Secondly, by denoting:

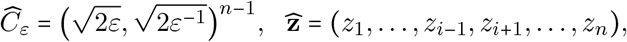

and:

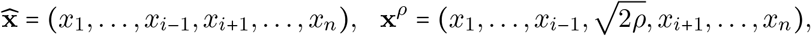

for *ρ* ≥ 0, it follows from (16) that:

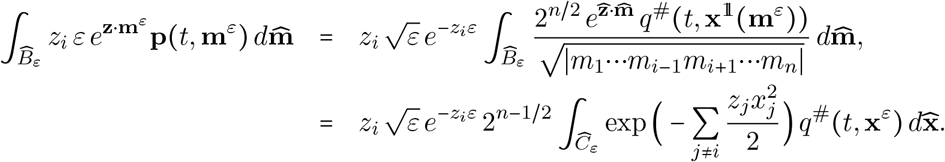

Since the continuous function *q*^#^(*t*, ·) is bounded in ℝ^*n*^ by (38) and (48), and since 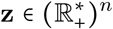, one then gets that:

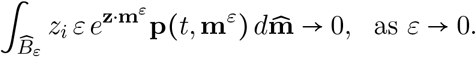

Similarly, we prove that:

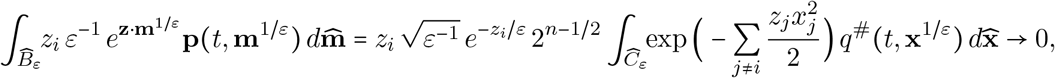

as *ε* → 0. Thirdly, from the computations done in the proof of Theorem 2.4, we already know that, for all 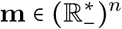:

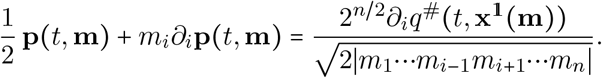

Hence, we have:

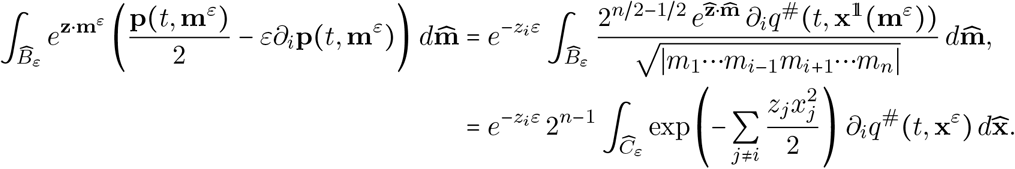

Since the function *∂_i_q*^#^(*t*, ·) is continuous and bounded in ℝ^*n*^ by (38) and (48), since *∂_i_q*^#^ (*t*, **x**^0^) = 0, by #-symmetry of *q*^#^ (*t*,·), and since 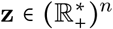, one then infers from Lebesgue’s dominated convergence theorem that:

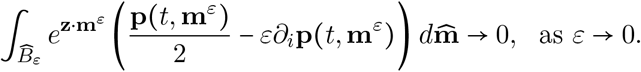

Furthermore, the integral:

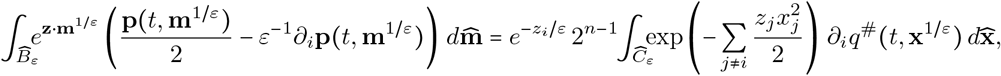

converges to 0 as *ε* → 0. Coming back to (58) and passing to the limit as *ε* → 0, it follows from the previous estimates that:

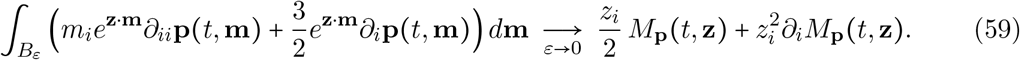

Let us finally remember (57) and that the functions:

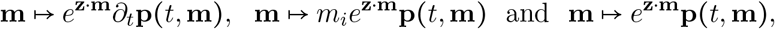

are in 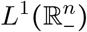 with integralsgiven by (56), (52) and (21), respectively. Together with (58) and (59), one concludes that:

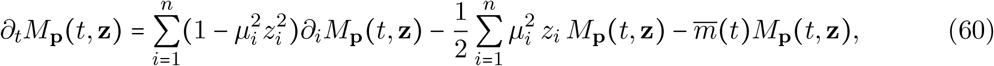

for every 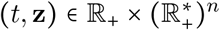. Since the right-hand side of the above equation is continuous in 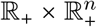, one infers that the function *∂_t_M*_**p**_ is extendable by continuity in 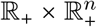 and (60) holds in 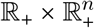. Owing to the definition *C*_p_ = log*M*_**p**_, one concludes that *∂_t_C*_**p**_ is continuous in 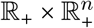 (finally, *C*_p_ is of class 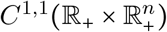) and:

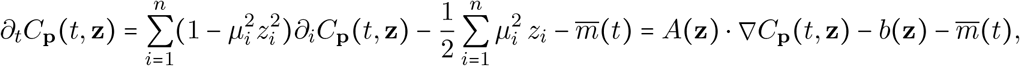

for all 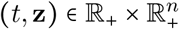, where *A* and *b* are as in (24). Therefore, (23) holds in 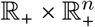 and the proof of Theorem 2.6 is thereby complete.

Before going into the proof of the remaining results, let us first observe that, in (23)–(24), 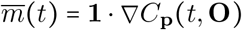, with:

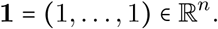

It turns out that, if *A*(**z**) = **1**, then equation (23) can be solved explicitely by the method of characteristics, as the following lemma shows (this lemma is used later in the proof of Proposition 2.8 in the general case *A*(**z**) given in (24)).

#### Lemma 4.3.

The Cauchy problem:

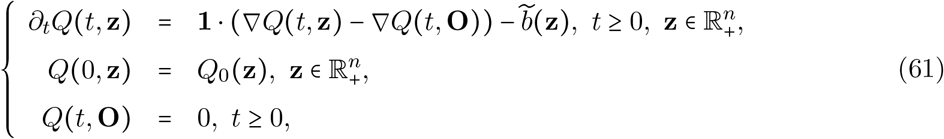

with 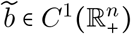 and 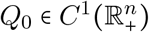 such that 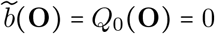, admits a unique 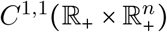 solution, which is given by the expression:

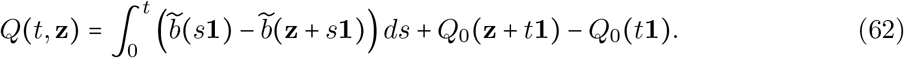

*Proof*. First of all, it is immediate to check that the function *Q* given by (62) is a 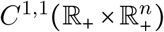 solution of (61). Let now *Q*_1_ and *Q*_2_ be two 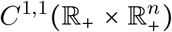 solutions of (61) and denote *Q* = *Q*_1_ − *Q*_2_. The function *Q* is of class 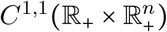 and obeys:

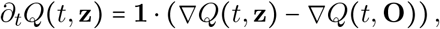

for all 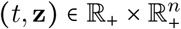, together with *Q*(0, **z**) = 0 for all 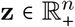 and *Q*(*t*, **O**) = 0 for all *t* ≥ 0. It remains to show that *Q* = 0 in 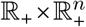. Fix any 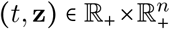. If *t* = 0, then *Q*(0, **z**) = 0, so let us assume that *t* > 0. Consider the *C*^1^([0,*t*]) function R defined by *R*(*s*) =*Q*(*t–s*, **z**+*s***1**)−*Q*(*t–s*, *s***1**) for *s* ϵ [0, *t*] (which is well defined since 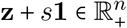). It follows from the equation satisfied by *Q* that, for all *s* ϵ [0,*t*], there holds:

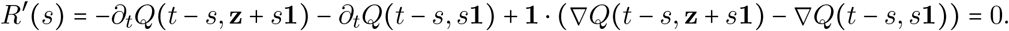

Hence, 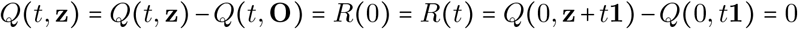, which is the desired conclusion.

#### Proof of Proposition 2.8.

In order to derive a general formula for the 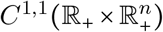 solution *C*_**p**_ of (23), we make a substitution of the spatial variable and use the previous special case described in Lemma 4.3. To do so, we set, for *t* ≥ 0 and 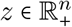,

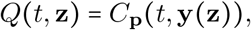

where **y**(**z**) = (*y*_1_(**z**),…, *y_n_*(**z**)) and *y_i_*(**z**) =tanh(*μ_i_z_i_*)/*μ_i_* for every 1 ≤ *i* ≤ *n*. Notice that 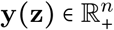 for every 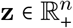. The function *Q* is of class 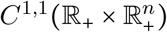 and:

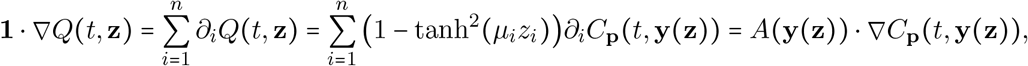

for all 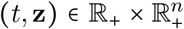, where *A* is given in (24). As 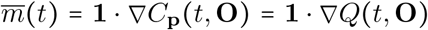 and *Q*(*t*, **O**) = *C*_**p**_(*t*, **O**) =log *M*_**p**_(*t*, **O**) = 0 by (15) and (21), it follows from (23) that:

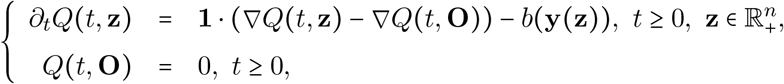

and *Q*(0, **z**) =*C*_**p**0_(**y**(**z**)) for all 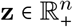. The functions *C*_**p**0_ ◦ **y** and 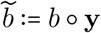 are of class 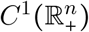 and 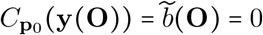. Therefore, Lemma 4.3 implies that:

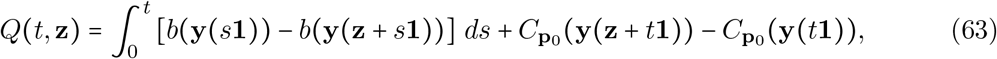

for all 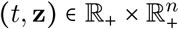. Consider now any *t* ϵ ℝ_+_ and **z** = (*z*_1_,…, *z_n_*) ϵ [0,1/*μ*_1_) × … × [0,1/*μ_n_*). Set:

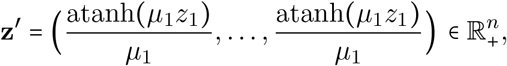

and observe that **y**(**z**′) =**z**. Hence, we have:

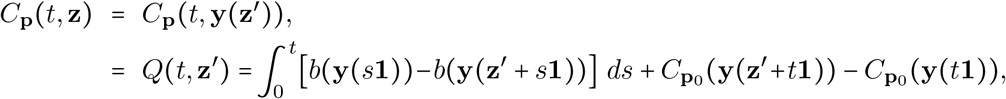

which leads straightforwardly to the formulae (26)–(27). Furthermore, for every *t* ϵℝ_+_, the formula 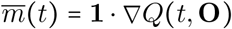 together with (63) easily yields (28). The proof of Proposition 2.8 is thereby complete.

#### Proof of Corollary2.7.

Let *p* be the fitness distribution, that is, *p*(*t,m*) *dm* is the pushforward measure of *q*(*t*, **x**) *d***x** by the map **x** ↦ − ‖**x**‖^2^/2. Let:

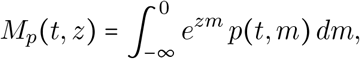

be the moment generating function of *p*. As the fitness *m* ϵℝ_−_ is the sum of the fitness components 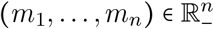, we have:

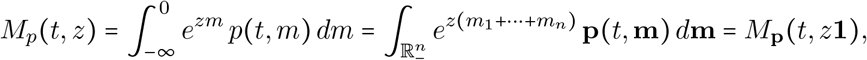

for all (*t, z*) ϵℝ_+_×ℝ_+_. This implies that *C_p_*(*t,z*) =*C*_**p**_(*t, z***1**) for all (*t, z*) ϵℝ_+_×ℝ_+_ and that *C_p_* is of class *C*^1,1^(ℝ_+_×ℝ_+_), with initial condition *C_p_*(0, ·) =*C*_**p**0_(·**1**). Thanks to the equations (23)–(24) satisfied by *C***p**, it follows that:

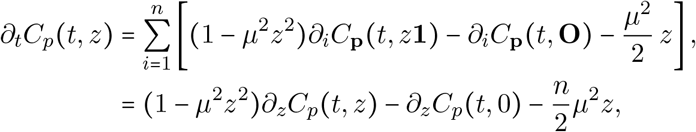

for all (*t, z*) ϵℝ_+_×ℝ_+_. This is the desired result and the proof of Corollary 2.7 is thereby complete.

#### Proof of Corollary 2.9.

We have seen in the proof of Corollary 2.7 that *C_p_*(*t, z*) =*C*_**p**_(*t, z***1**) for all (*t, z*) ϵℝ_+_×ℝ_+_. Thus, formulae (26)–(28) straightforwardly yield (29)–(30) for *t* ≥ 0 and *z* ϵ [0,1/*μ*).

### 4.4 Stationary states

#### Proof of Theorem 2.10.

We use the notations of Proposition 2.8. Let **z** ϵ [0,1/*μ*_1_) ×…× [0,1/*μ_n_*). As 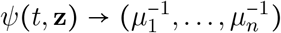 as *t* →+∞, the continuity of *C*_**p**0_ and the formulae (26)–(27) yield that:

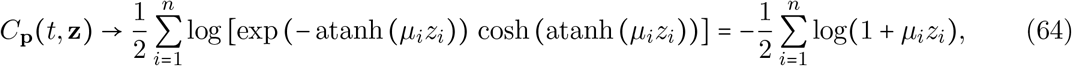

as *t* →+∞. It then follows from the generalization of the Curtiss theorem [46] that, if the limit as *t* →+∞of the cumulant generating functions *C*****p****(*t*, ·) is the cumulant generating function of **p**_∞_ given by (31) in some subset of 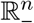 with non-empty interior, then the distributions **p**(*t*, ·) weakly converge to **p**_∞_ as *t* →+∞. So let us compute the CGF of **p**_∞_. For all 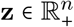, Fubini’s theorem yields that:

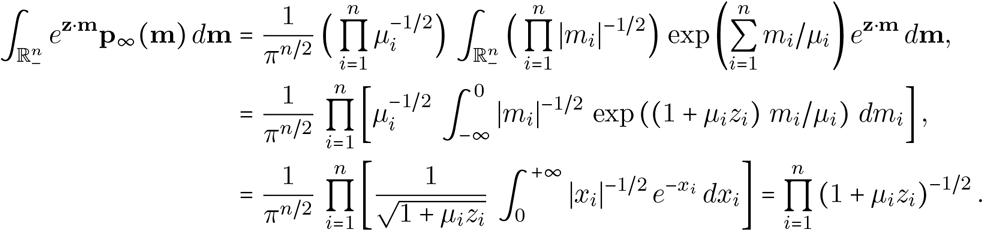

Hence, the CGF of **p**_∞_ is equal to the function 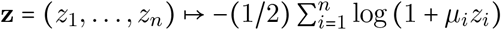, that is, the limit in (64). As a consequence, the distributions **p**(*t*, ·) weakly converge to **p**_∞_ in 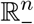 as *t* →+∞.

On the other hand, thanks to Proposition 2.8, we also know that:

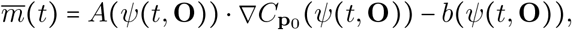

for every *t* ≥ 0, with 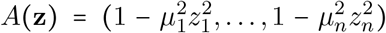 and 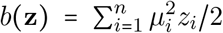. Notice that 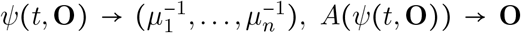 and 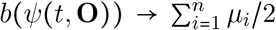, as *t* →+∞. Hence, 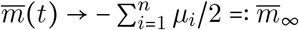, as *t* →+∞. It is also straightforward to check that:

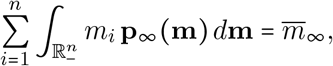

and that **p**_∞_ is a classical 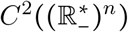 solution of (32) (this property is also a consequence of the fact that **p** satisfies (17) and the distributions **p**(*t*, ·) weakly converge to **p**_∞_ as *t* →+∞). The proof of Theorem 2.10 is thereby complete.

#### Proof of Corollary 2.11.

By the same arguments as in the proof of Theorem 2.10, thanks to (29), we see that, for all *z* ϵ [0,1/*μ*), *C_p_*(*t, z*) → − (1/2) log(1 + *μz*) as *t* →+∞. This limiting function corresponds to the cumulant generating function of a random variable distributed according to −Γ(*n*/2,*μ*). Since a distribution is uniquely determined by its cumulant generating function, this implies that *p*_∞_ is the probability density function of this random variable, *i.e*., for all *m* < 0,

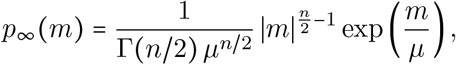

where 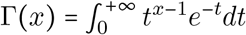 is the standard Gamma function.

#### Proof of Corollary 2.12.

We assume here that *q*_0_ is #-symmetric, that is, 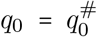. As already emphasized in Section 2.1, the uniqueness for problem (10) implies that *q*(*t*, ·) is also #-symmetric at each time *t* ≥ 0. Proposition 2.3 (or formula (16)) yields:

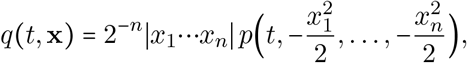

for all (*t*, **x**) ϵ ℝ_+_× (ℝ*)^*n*^ and, for each function 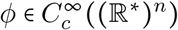:

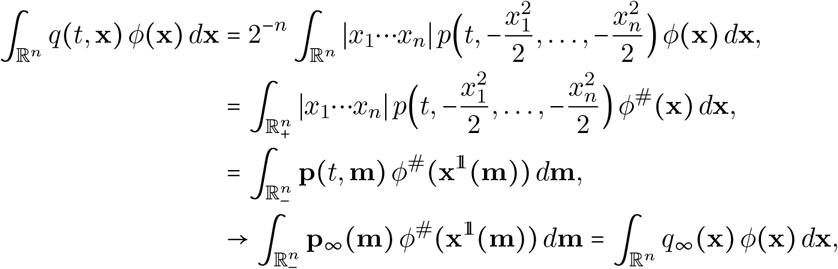

as *t* → + ∞, with 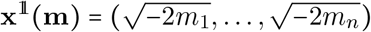 and:

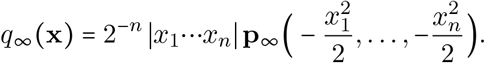

The above formula corresponds to (33). Furthermore, since 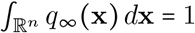 and since 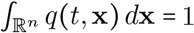, for every *t* ≥ 0, it then easily follows from the previous estimates that:

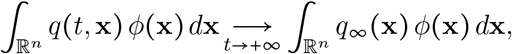

for every 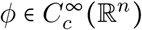. In other words, the distributions *q*(*t*, ·) weakly converge in ℝ^*n*^ to *q*_∞_ as *t* →+∞. Lastly, the formula 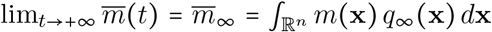 is a consequence of the previous arguments and Theorem 2.10. The proof of Corollary 2.12 is thereby complete.

### 4.5 Plateaus: proofs of Proposition 2.13 and Remark 2.14

#### Proof of Proposition 2.13.

We show in this section that, given an initial phenotype **x**_0_ = (*x*_0,1_,…, *x*_0,*n*_) ϵ ℝ^*n*^, a value *μ*_1_ > 0, and a duration *T* > 0, we can choose some positive real numbers *μ*_2_,…, *μ_n_* (here, *n* ≥ 2) such that the mean fitness:

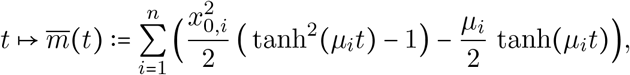

is close to each of the plateaus:

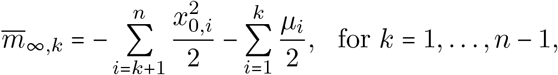

at least during a time interval of duration *T*. We recall that the above formula for 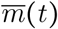 corresponds to the limit of formula (28), when the initial conditions *q*_0_ approach the Dirac distribution *δ*_xo_.

More precisely, we are given **x**_0_ = (*x*_0,1_,…, *x*_0,*n*_) ϵ ℝ^*n*^, *T* > 0, *ε* > 0 and *μ*_1_ > 0, and we shall prove the existence of 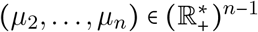 such that, for each *k* ϵ 〚 1,*n* − 1〛, the set:

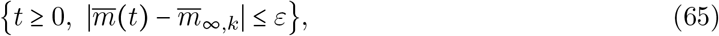

contains an interval of length at least equal to *T*. In that respect, we firstly define some functions:

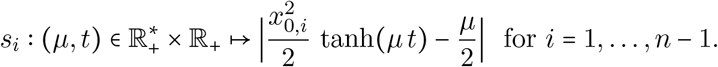

Secondly, by iteration for *k* = 1,…, *n* − 1, we can then define:

◊ a function 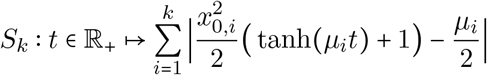,
◊ a time *τ_k_* > *τ*_*k* − 1_ + *T*, (with *τ*_0_ = − *T*) such that:

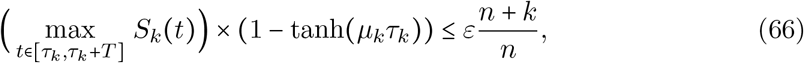
◊ a real number *μ*_*k*+ 1_ ϵ (0, *μ_k_*) such that:

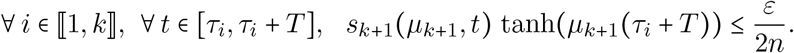

Note that the last property implies that:

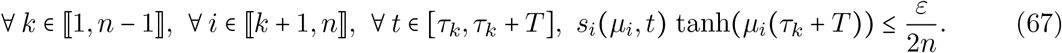

Fix now an index *k* ϵ 〚 1, *n* − 1〛 and a time 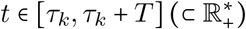. There holds:

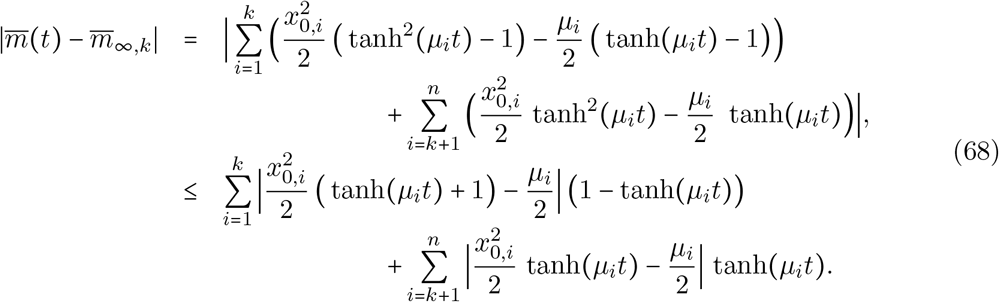

As *t* ϵ [*τ_k_,τ_k_*+*T*], we have 0 < 1 − tanh(*μ_i_t*) ≤ 1 − tanh(*μ_k_τ_k_*) for every *i* ϵ 〚 1, *k*〛 (remember that 0 < *μ_k_* < *μ*_*k* − 1_ < … < *μ*_1_), whereas tanh(*μ_i_t*) ≤ tanh(*μ_i_*(*τ_k_*+*T*)) for every *i* ϵ 〚*k* + 1,*n*〛. It then follows from (66)–(68) that, for every *k* ϵ 〚 1,*n* − 1〛 and *t* ϵ [*τ_k_,τ_k_*+*T*]:

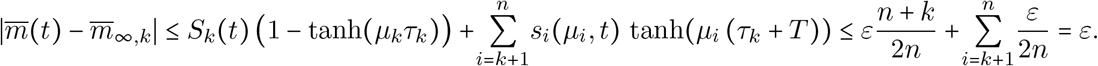

Thus, with this choice of 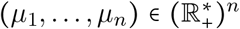, each set defined in (65) contains an interval of length at least equal to *T*. This proves Proposition 2.13.

#### Proof of results in Remark 2:14.

Take *n* =2, *μ*_1_ = 1, *μ*_2_ = 10^−*k*^ for some *k* ≥ 1 and *x*_0,1_ = *x*_0,2_ = 1. Differentiating the expression 34, we observe that:

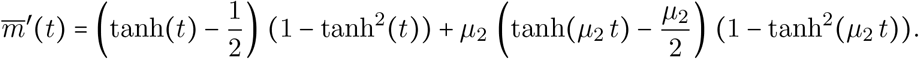

We observe that, as 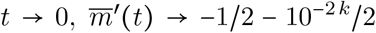. Then 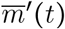 rapidly increases, to reach significantly positive values, e.g., at 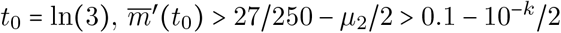.

Then, consider the interval:

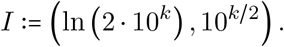

For each *t* in *I*, 1 − tanh^2^(*t*) <4 exp(−21) < 10^−2*k*^ and tanh(*μ*_2_*t*) − *μ*_2_/2 < *μ*_2_*t* < 10^*k*/2^. Thus,

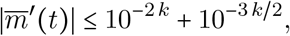

which means that 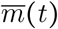 remains stable within this interval, corresponding to a part of the plateau. Latter on, 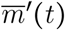 reaches again significantly positive values which are significantly higher than in this interval, e.g., at 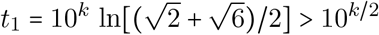, straightforward computations show that:

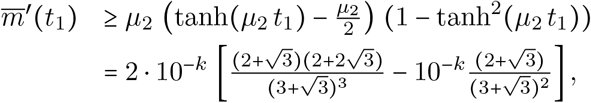

which show that 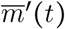 is of order 10^−*k*^ at *t*_1_, vs 10^−3*k*/2^ in the interval I.

## A A formal derivation of the diffusive approximation of the mutation effects

The goal of this appendix is to give a formal justification of the diffusion term in (4). The case *n* = 1 is classical and can be found *e.g*. in [37, 43]. The anisotropic case *n* ≥ 2 is less standard, but it will easily follow from the same arguments.

Namely, we assume that the mutation effects on phenotypes follow a normal distribution 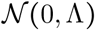, with Λ=diag(λ_1_,…,λ_*n*_) and λ_*i*_ > 0 for each *i* ϵ 〚 1,*n*〛, and that these mutations occur with a rate *U* > 0. In other words, the dynamics of the phenotype distribution under the mutation effects only (*i.e*., without selection) can be described by an integro-differential equation:

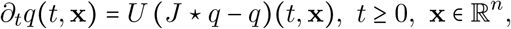

where * is the standard convolution product in ℝ^*n*^ defined by:

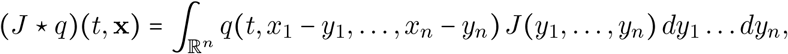

and *J* the (Gaussian) probability density function associated with the normal distribution 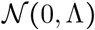.

Formally, by writing a Taylor expansion of *q*(*t*, **x** − **y**) at **x**:

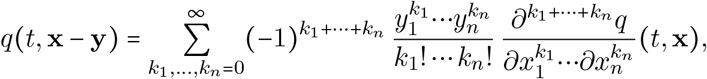

and by defining the central moments of the normal distribution:

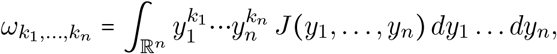

we then get that:

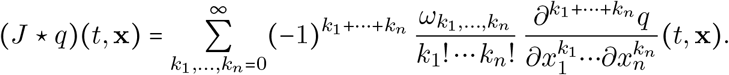

Since *ω*_*k*_1_,…,*k*_*n*__ = 0 if at least one of the *k_i_*’s is odd, and since:

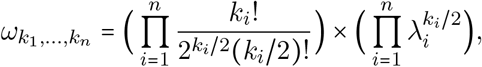

otherwise, one infers in particular that the second-order moments with even indexes are such that *ω*_0_,…,0,*k_i_*=2,0,…,0=λ_*i*_. Assuming that 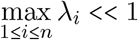, we may formally neglect the moments of *k*_1_ + … +*k_n_* ≥ 4, leading to:

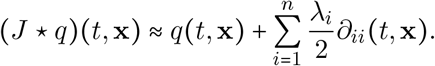

Finally, setting 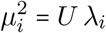, we obtain:

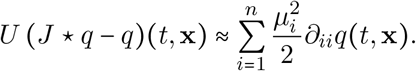

## Acknowledgments

This work was supported by the French Agence Nationale de la Recherche (ANR-13-ADAP-0016 “Silentadapt” to G.M., ANR-13-ADAP-0006 “MeCC” to L.R., ANR-14-CE25-0013 “NON-LOCAL” to F.H. and L.R. and ANR-18-CE45-0019 “RESISTE” to F.H., F.L., G.M. and L.R.). This work was fostered by stimulating discussions with Thomas Lenormand. The authors also thank the reviewers for valuable comments and suggestions.

## Footnotes

1. This means that the partial derivatives of *C*_**p**_ with respect to the variables *t* and **z** exist and are continuous in 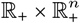.

## References

[1] M. Alfaro and R. Carles, Explicit solutions for replicator-mutator equations: Extinction versus acceleration, SIAM Journal on Applied Mathematics 74 (2014) 1919–1934.

[2] M. Alfaro and R. Carles, Replicator-mutator equations with quadratic fitness, Proceedings of the American Mathematical Society 145 (2017) 5315–5327.

[3] M. Alfaro and M. Veruete, Evolutionary branching via replicator-mutator equations, Journal of Dynamics and Differential Equations (2018) 1–24.

[4] D. G. Aronson and P. Besala, Parabolic equations with unbounded coefficients, Journal of Differential Equations 3 (1967) 1–14.

[5] E. Baake, A. G. Casanova, S. Probst and A. Wakolbinger, Modelling and simulating Lenski’s long-term evolution experiment, Theoretical population biology 127 (2019) 58–74.

[6] G. Barles, S. Mirrahimi and B. Perthame, Concentration in Lotka-Volterra parabolic or integral equations: a general convergence result, Methods and Applications of Analysis 16 (2009) 321–340.

[7] A. G. Casanova, N. Kurt, A. Wakolbinger and L. Yuan, An individual-based model for the Lenski experiment, and the deceleration of the relative fitness, Stochastic Processes and their Applications 126 (2016) 2211–2252.

[8] N. Champagnat, R. Ferrière and S. Méléard, Unifying evolutionary dynamics: from individual stochastic processes to macroscopic models, Theoretical population biology 69 (2006) 297–321.

[9] V. A. de Crécy-Lagard, J. Bellalou, R. Mutzel and P. Marlière, Long term adaptation of a microbial population to a permanent metabolic constraint: overcoming thymineless death by experimental evolution of *Escherichia coli*, BMC biotechnology 1 (2001) 10.

[10] O. Diekmann, P.-E. Jabin, S. Mischler and B. Perthame, The dynamics of adaptation: an illuminating example and a Hamilton–Jacobi approach, Theoretical population biology 67 (2005) 257–271.

[11] S. Elena and R. Sanjuán, Climb every mountain?, Science 302 (2003) 2074–2075.

[12] S. S. Fong and B. Ø. Palsson, Metabolic gene-deletion strains of *Escherichia coli* evolve to computationally predicted growth phenotypes, Nature genetics 36 (2004) 1056.

[13] C. Fraïsse and J. J. Welch, The distribution of epistasis on simple fitness landscapes, Biology letters 15 (2019) 20180881.

[14] A. Friedman, Partial Differential Equations of Parabolic Type (Prentice-Hall, Englewood Cliffs, NJ, 1964).

[15] S. Gandon and S. Mirrahimi, A Hamilton–Jacobi method to describe the evolutionary equilibria in heterogeneous environments and with non-vanishing effects of mutations, Comptes Rendus Mathematique 355 (2017) 155–160.

[16] P. J. Gerrish and P. D. Sniegowski, Real time forecasting of near-future evolution, Journal of the Royal Society Interface 9 (2012) 2268–2278.

[17] M.-E. Gil, F. Hamel, G. Martin and L. Roques, Mathematical properties of a class of integro-differential models from population genetics, SIAM J Appl Math 77 (2017) 15361561.

[18] M.-E. Gil, F. Hamel, G. Martin and L. Roques, Dynamics of fitness distributions in the presence of a phenotypic optimum: an integro-differential approach, Nonlinearity (2019) in press.

[19] B. H. Good and M. M. Desai, The impact of macroscopic epistasis on long-term evolutionary dynamics, Genetics 85 (2015) 177–190.

[20] P. D. Hislop and I. M. Sigal, Introduction to spectral theory: With applications to Schrödinger operators, volume 113 (Springer Science & Business Media, 2012).

[21] S. Kryazhimskiy, G. Tkačik and J. B. Plotkin, The dynamics of adaptation on correlated fitness landscapes, Proceedings of the National Academy of Sciences 106 (2009) 18638–18643.

[22] R. A. LaCroix, t. E. Sandberg, E. J. O’Brien, J. Utrilla, A. Ebrahim, G. I. Guzman, R. Szubin, B. O. Palsson and A. M. Feist, Use of adaptive laboratory evolution to discover key mutations enabling rapid growth of *Escherichia coli* K-12 MG1655 on glucose minimal medium, Applied and Environmental Microbiology 81 (2015) 17–30.

[23] R. E. Lenski, M. R. Rose, S. C. Simpson and S. C. Tadler, Experimental evolution in *Escherichia coli.* I. Adaptation and divergence during 2,000 generations, The American Naturalist 138 (1991) 1315–1341.

[24] R. E. Lenski and M. Travisano, Dynamics of adaptation and diversification: a 10,000-generation experiment with bacterial populations, Proceedings of the National Academy of Sciences 91 (1994) 6808–6814.

[25] A. Lorz, S. Mirrahimi and B. Perthame, Dirac mass dynamics in multidimensional nonlocal parabolic equations, Communications in Partial Differential Equations 36 (2011) 1071–1098.

[26] A. Lunardi, Schauder theorems for linear elliptic and parabolic problems with unbounded coefficients in ℝ^*n*^, Studia Mathematica 128 (1998) 171–198.

[27] G. Martin, Fisher’s geometrical model emerges as a property of complex integrated phenotypic networks, Genetics 197 (2014) 237–255.

[28] G. Martin, S. F. Elena and T. Lenormand, Distributions of epistasis in microbes fit predictions from a fitness landscape model, Nature Genetics 39 (2007) 555.

[29] G. Martin and T. Lenormand, A general multivariate extension of Fisher’s geometrical model and the distribution of mutation fitness effects across species, Evolution 60 (2006) 893–907.

[30] G. Martin and T. Lenormand, The fitness effect of mutations across environments: a survey in light of fitness landscape models, Evolution 60 (2006) 2413–2427.

[31] G. Martin and T. Lenormand, The fitness effect of mutations across environments: Fisher’s geometrical model with multiple optima, Evolution 69 (2015) 1433–1447.

[32] G. Martin and L. Roques, The non-stationary dynamics of fitness distributions: Asexual model with epistasis and standing variation, Genetics 204 (2016) 1541–1558.

[33] I. S. Novella, E. A. Duarte, S. F. Elena, A. Moya, E. Domingo and J. J. Holland, Exponential increases of RNA virus fitness during large population transmissions, Proceedings of the National Academy of Sciences 92 (1995) 5841–5844.

[34] L. Perfeito, A. Sousa, T. Bataillon and I. Gordo, Rates of fitness decline and rebound suggest pervasive epistasis, Evolution 68 (2014) 150–162.

[35] B. Perthame and G. Barles, Dirac concentrations in Lotka-Volterra parabolic PDEs, Indiana University Mathematics Journal (2008) 3275–3301.

[36] M. H. Protter and H. F. Weinberger, Maximum Principles in Differential Equations (Prentice-Hall, Englewood Cliffs, NJ, 1967).

[37] L. Roques, Modèles de réaction-diffusion pour l’écologie spatiale (Editions Quae, 2013).

[38] F. Rosenzweig and G. Sherlock, Experimental evolution: prospects and challenges, Genomics 104 (2014) v.

[39] S. Schoustra, S. Hwang, J. Krug and J. A. G. de Visser, Diminishing-returns epistasis among random beneficial mutations in a multicellular fungus, Proceedings of the Royal Society B: Biological Sciences 283 (2016) 20161376.

[40] P. D. Sniegowski and P. J. Gerrish, Beneficial mutations and the dynamics of adaptation in asexual populations, Philosophical Transactions of the Royal Society B: Biological Sciences 365 (2010) 1255–1263.

[41] O. Tenaillon, The utility of Fisher’s geometric model in evolutionary genetics, Annual Review of Ecology, Evolution, and Systematics 45 (2014) 179–201.

[42] L. S. Tsimring, H. Levine and D. A. Kessler, RNA virus evolution via a fitness-space model, Physical review letters 76 (1996) 4440–4443.

[43] P. Turchin, Quantitative Analysis of Movement: Measuring and Modeling Population Redistribution in Animals and Plants (Sinauer, Sunderland, MA, 1998).

[44] D. Waxman and J. R. Peck, Pleiotropy and the preservation of perfection, Science 279 (1998) 1210–1213.

[45] M. J. Wiser, N. Ribeck and R. E. Lenski, Long-term dynamics of adaptation in asexual populations, Science (2013) 1243357.

[46] A. L. Yakymiv, A generalization of the Curtiss theorem for moment generating functions, Math. Notes 90 (2011) 920–924.

